# BET Inhibitors Target the SCLC-N subtype Small Cell Lung Cancer by Blocking NEUROD1 Transactivation

**DOI:** 10.1101/2021.10.25.465771

**Authors:** Haobin Chen, Lisa Gesumaria, Young-Kwon Park, Trudy G. Oliver, Dinah S. Singer, Kai Ge, David S. Shrump

## Abstract

Small cell lung cancer (SCLC) is a recalcitrant malignancy that urgently needs new therapies. Four master transcription factors (ASCL1, NEUROD1, POU2F3, and YAP1) are identified in SCLC, and each defines the transcriptome landscape of one molecular subtype. These master factors have not been directly druggable, and targeting their transcriptional coactivator(s) could provide an alternative approach. Here, we identify that BET bromodomain proteins physically interact with NEUROD1 and function as its transcriptional coactivators. Using CRISPR knockout and ChIP-seq, we demonstrate that NEUROD1 plays a critical role in defining the landscapes of BET bromodomain proteins in the SCLC genome. Targeting BET bromodomain proteins by BET inhibitors leads to broad suppression of the NEUROD1-target genes, especially those associated with superenhancers, and reduces SCLC growth *in vitro* and *in vivo*. LSAMP, a membrane protein in the IgLON family, was identified as one of the NEUROD1-target genes mediating BET inhibitor sensitivity in SCLC. Altogether, our study reveals that targeting transcriptional coactivators could be a novel approach to blocking the master transcription factors in SCLC for therapeutic purposes.

**Significance:** Small cell lung cancer (SCLC) is the most aggressive form of lung malignancies, and little progress has been made to improve its outcome in the past two decades. It is now recognized that SCLC is not a single disease but has at least four molecular subtypes, and each subtype features the expression of one master transcription factor. Unfortunately, these master transcription factors are not directly druggable. Here, we identified BET bromodomain proteins as the transcriptional coactivators of NEUROD1, one of the master transcription factors in SCLC. Blocking BET bromodomain proteins with inhibitors suppresses NEUROD1-target genes and reduces tumor growth. Our results demonstrate that blocking transcriptional coactivators could be an alternative approach to targeting the master transcription factors in SCLC.

## Introduction

Small cell lung cancer (SCLC) is a recalcitrant malignancy with few treatment options (1). Despite recent advances in therapeutics, durable responses are typically infrequent in extensive-stage SCLC, and its prognosis remains dismal (2, 3). Growing evidence has shown that SCLC is a heterogeneous group of cancers, and a recently proposed classification divided it into four molecular subtypes (SCLC-A, SCLC-N, SCLC-P, and SCLC-Y) according to the expression of various master transcription factors (TFs; ASCL1, NEUROD1, POU2F3, and YAP1, respectively) (4, 5). While there is still debate whether an SCLC-Y subtype tumor truly exists, several studies have confirmed the presence of other three subtypes of tumors (6, 7). Emerging evidence suggests that various molecular subtypes of SCLC have different therapeutic susceptibilities, likely because of their distinct transcriptome profiles dictated by the master TFs (8, 9). There have been substantial interests in developing precision therapies by targeting individual subtypes of SCLC (8, 10), but these master TFs have not been found druggable.

NEUROD1 is a neurogenic basic helix-loop-helix TF and plays a critical role in neuronal differentiation (11). Previous studies in embryonic stem cells and microglia cells found that NEUROD1 could induce the neuronal genes by directly binding to the target heterochromatic promoters and enhancers, leading to chromatin remodeling and TF reprogramming (12, 13). NEUROD1 promotes cell migration and invasion of SCLC through NTRK2 (previously known as TrkB) and NCAM (14). Although both NEUROD1 and ASCL1 regulate neuroendocrine genes in SCLC, they occupy primarily distinct genomic regions except for 5-7% overlaps (15). Little is known how NEUROD1 activates its downstream gene transcription and what genes, and to what degree, are dependent on NEUROD1 in SCLC. Such information is critical for developing new strategies to target this master TF.

The bromodomain and extraterminal domain (BET) family of proteins, including BRD2, BRD3, BRD4, and BRDT, are involved in multiple aspects of transcriptional regulation (16). Two bromodomains (BDs) at the N-terminus of this family of proteins are essential for binding to active chromatin (16, 17). BET inhibitors (BETis) selectively bind to these BDs and block BET bromodomain proteins from accessing active chromatin, resulting in the suppression of gene transcription (18, 19). Although BET bromodomain proteins are broadly associated with gene promoters and enhancers, BETi only inhibits a subset of genes, particularly those with unusually high levels of BRD4 occupancy at their enhancer sites known as superenhancers (SEs) (20, 21).

We hypothesized that targeting the transcriptional cofactors of these master TFs could effectively block their transcriptional activity. Here we demonstrated that BET bromodomain proteins function as the transcriptional coactivators of NEUROD1 and could be a therapeutic target in SCLC-N subtype tumors.

## Results

### NEUROD1 is essential for the growth and tumorigenicity of SCLC-N cell lines

To determine whether NEUROD1 is essential for SCLC-N tumors, we used CRISPR to knock out this gene in H446 cells (an SCLC-N cell line) with two distinct guide RNAs (gRNAs). Western blots confirmed a complete loss of NEUROD1 protein expression in the knockout (KO) clones of both gRNAs (Figs. 1A and S1A). We validated NEUROD1 KO by identifying indels in the targeted NEUROD1 genomic sequences (Fig. S1B-C) and finding a drastically decreased expression of two known NEUROD1-target genes - *NEUROD2* and *NHLH1* (Figs. 1B and S1D).

**Figure 1.**
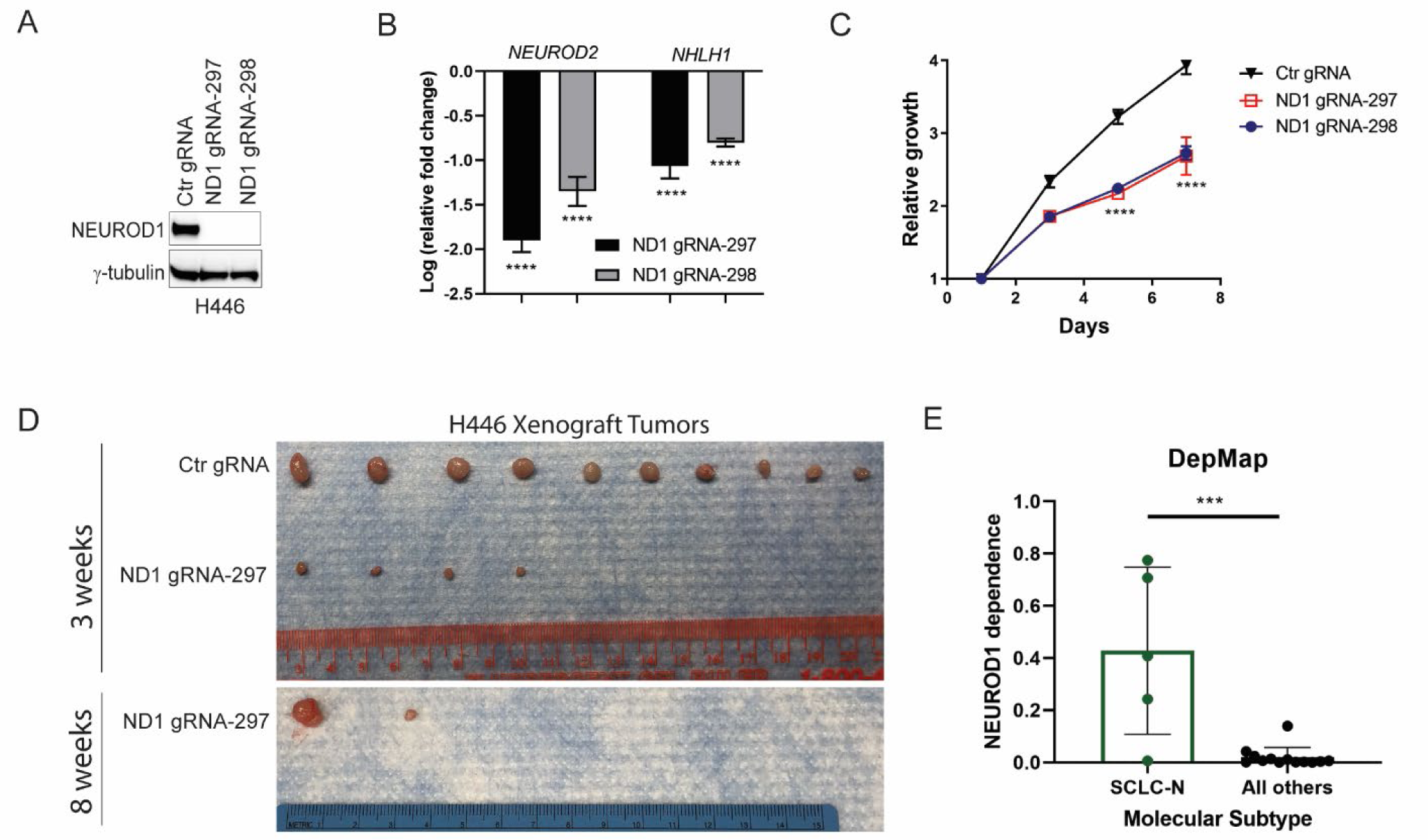
NEUROD1 is essential for the growth and tumorigenicity of the SCLC-N cell lines. **A)** Western blots showing a complete loss of NEUROD1 protein expression in two H446 NEUROD1-KO clones. One representative clone established from each gRNA (#297 cl. 15 and #298 cl. S3) was shown. **B)** Expression of two NEUROD1-target genes, *NEUROD2* and *NHLH1,* in the H446 NEUROD1-KO cells (#297 cl. 15 and #298 cl. S3). The gene expression was measured by qRT-PCR and plotted relative to control cells. **C)** The H446 NEUROD1-KO cells (#297 cl. 15 and #298 cl. S3) proliferated substantially slower than control cells. Cell numbers were indirectly measured using the CellTiter Glo 2.0 assay every other day, and the results were compared to the reading on day one. Error bar represents SD of 4 replicates. **D)** NEUROD1 KO impaired the tumorigenicity of H446 cells. Top, the xenografts formed three weeks after injecting 1x10^6^ H446 control and NEUROD1-KO cells (#297 cl. 15) into the flanks of nude mice (n=10 per group). Bottom, xenografts formed eight weeks after injection of 1x10^6^ NEUROD1-KO cells (#297 cl. 12; n=10). **E)** Comparing *NEUROD1* gene dependency between the SCLC-N and other subtypes of SCLC lines, using the data extracted from the DepMap project (Achilles Gene Dependency – CRISPR, Version 2020, Q2). The significance of two-group comparisons was determined by the ANOVA test with Dunnett’s multiple test correction (B, C) and the Student’s *t*-test (E). ***, P < 0.001; ****, P < 0.0001. Ctr, Control; gRNA, guide RNA; ND1, NEUROD1.

We next assessed the impact of NEUROD1 KO on the growth and tumorigenicity of H446 cells. As shown in Figure 1C, KO of NEUROD1 rendered H446 cells to proliferate much slower than control cells. Furthermore, when the KO cells were injected subcutaneously into athymic mice, fewer and smaller xenografts formed even after five additional weeks of growth (Fig. 1D). Consistent with these findings, the CRISPR screens from the Dependency Map (DepMap) project showed NEUROD1 is a high dependency gene in the SCLC-N lines (Fig. 1E). Collectively, these results demonstrate that NEUROD1 is essential for the growth and tumorigenicity of SCLC-N cell lines.

### BET bromodomain proteins interact with NEUROD1

We hypothesized that targeting transcriptional coactivator(s) can block the transactivation of NEUROD1. To identify transcriptional factors involved in NEUROD1 transactivation, we first performed ChIP-sequencing (ChIP-seq) to map NEUROD1 genome occupancy in the H446 control and KO cells. Enrichment of the NEUROD1/NEUROG2 binding motif was found, as expected, in the genomic sequences corresponding to NEUROD1 peaks (Fig. S2). As shown in Figure 2A, most of the NEUROD1 peaks locate in intronic (45.8%) and intergenic regions (45.9%), and the remaining was present at transcription start sites (TSS; 1.12%), promoter-TSS sites (3.88%), and exons + 3’ or 5’ untranslated regions (UTRs; 3.28%). A metagene analysis showed that NEUROD1 binds to the genomic regions within seven kilobases upstream of TSS and downstream of transcription end sites, suggesting that it binds to both promoters and distal regulatory elements, such as enhancers (Fig. 2B).

**Figure 2.**
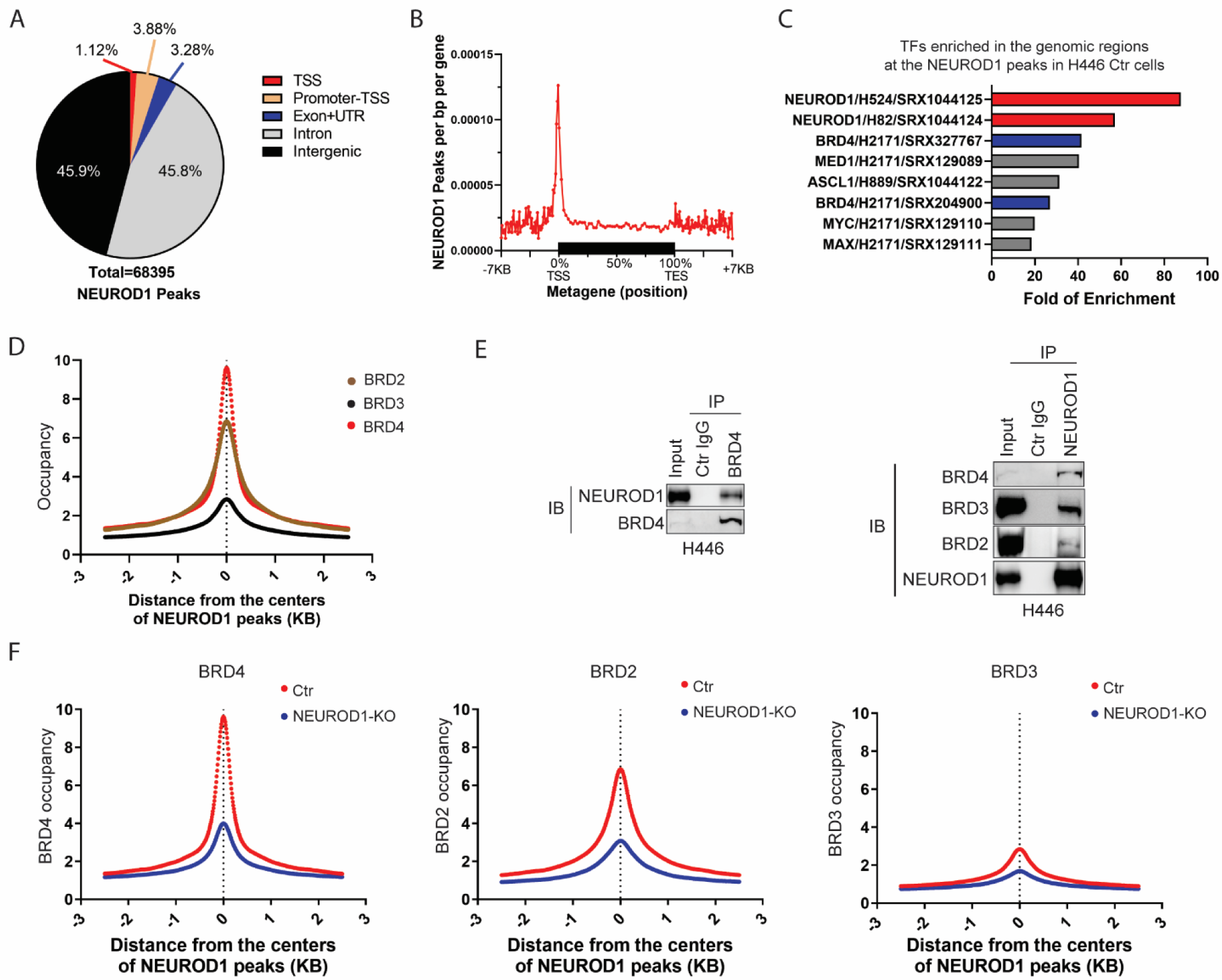
NEUROD1 interacts with BET bromodomain proteins and defines their genomic landscapes in SCLC. **A)** A breakdown of the genomic locations of the NEUROD1 peaks identified by ChIP-seq in H446 control cells (control gRNA). **B)** A metagene profile of NEUROD1 occupancy in H446 control cells. **C)** The bar graph shows the top 8 TF candidates identified by the Enrichment Analysis at ChIP-atlas.org that occupy the same genomic regions as NEUROD1 in H446 control cells. NEUROD1 and BRD4 were highlighted. **D)** BRD2, BRD3, and BRD4 occupancy near the centers of NEUROD1-occupied regions in H446 control cells (2.5 KB bilateral). **E)** Co-IP assays using H446 nuclear extracts. Left, detection of NEUROD1 after IP of BRD4. Right, detection of BRD2, BRD3, and BRD4 after IP of NEUROD1. **F)** Effect of NEUROD1 KO on BRD4, BRD2, and BRD3 occupancy near the centers of the NEUROD1- occupied regions in H446 control cells. IB, immunoblotting; IP, immunoprecipitation; TSS, transcription start site; TES, transcription end site; UTR, untranslated region.

To identify transcription factors enriched in the NEUROD1-occupied genomic regions, we performed an enrichment analysis (ChIP-atlas.org; ref. (22)) using the archived ChIP-seq datasets in lung cells. Among the eight identified candidates, the top two are NEUROD1 in SCLC-N cell lines - H524 and H82, demonstrating the validity of this approach (Fig. 2C). BRD4 and MED1, two markers of active enhancers, also showed high degrees of enrichment. ASCL1, known to occupy some genomic regions in common with NEUROD1 (15), was one of the candidates. We chose BRD4 for further study because there are existing inhibitors to target this gene and its association with NEUROD1 has not yet been characterized.

To confirm the colocalization of BRD4 and NEUROD1 in the SCLC genome, we mapped the genome occupancies of BRD4 and its two analogs - BRD2 and BRD3 in H446 cells using ChIP-seq. Figure 2D shows that all three BET family proteins colocalized with NEUROD1, with their occupancies summitting at the centers of the NEUROD1-occupied genomic regions. To determine whether NEUROD1 physically interacts with BET bromodomain proteins, we performed co-immunoprecipitation assays using the nuclear extracts from H446 and H524 cells. Figures 2E and S3 show that endogenous NEUROD1 immunoprecipitated with the BET family proteins and vice versa in both cell lines, indicating physical interaction between these proteins.

**Figure 3.**
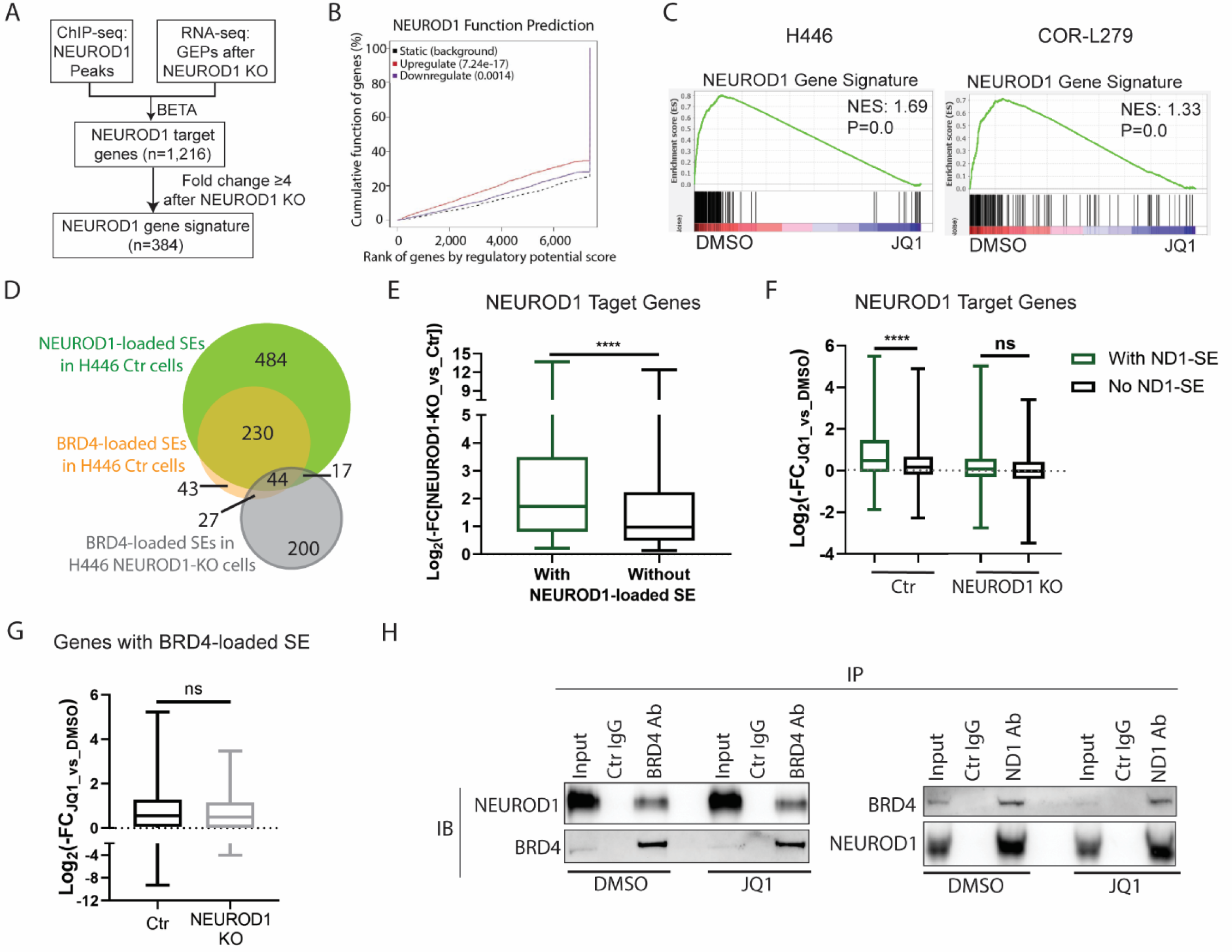
BETi suppresses NEUROD1-target genes, particularly those associated with superenhancers. **A)** A flow chart illustrating the process of identifying NEUROD1-target genes and selecting a subset of these genes to constitute a NEUROD1 gene signature. **B)** The BETA algorithm predicted that NEUROD1 primarily functions as a transcriptional activator. **C)** GSEA plots showing depletion of NEUROD1 gene signature by JQ1 in H446 (left; 1 µM, 24 hrs) and COR-L279 cells (right; 0.5 µM, 24 hrs). **D)** Venn diagrams showing overlaps between the NEUROD1- and BRD4-loaded SEs in H446 control and NEUROD1-KO cells. **E)** Comparing gene expression alterations between the NEUROD1-target genes with NEUROD1-loaded SEs and those without in H446 cells upon NEUROD1 KO. **F)** Differential effects of JQ1 (1µM, 24 hrs) on the expression of the NEUROD1-target genes with or without the NEUROD1-loaded SEs in H446 control (left two) and NEUROD1-KO cells (right two). **G)** JQ1 (1µM, 24 hrs) suppressed the genes with BRD4-loaded SEs to a similar degree in the H446 control and NEUROD1-KO cells. **H)** co-IP following JQ1 treatment (1µM, 6 hrs) in H446 cells. Left, detection of NEUROD1 after IP of BRD4. Right, detection of BRD4 after IP of NEUROD1. (E-G) Whiskers represent minimum to maximum, and boxes show the first quartile, median, and third quartile; the significance of two-group comparisons was determined using the Student’s *t*-test. ****, P<0.0001; ns: not significant. BETA, the Binding and Expression Target Analysis; FC, fold change; GEP, gene expression profile; IB, immunoblotting; IP, immunoprecipitation; NES, normalized enrichment score; SE; superenhancer.

We next assessed the impact of NEUROD1 KO on the genome occupancies of BET family proteins. As shown in Figure 2F, the occupancy of BET bromodomain proteins decreased substantially in the NEUROD1-occupied genomic regions upon NEUROD1 KO, suggesting that NEUROD1 directs the binding of BET bromodomain proteins in the SCLC genome. To determine whether these changes are limited to the NEUROD1-occupied genomic regions, we grouped all BRD4 peaks in H446 control cells on the basis of the presence or absence of NEUROD1 co-occupancy (Fig. S4, density plot 1 vs. 4). NEUROD1 KO abrogated BRD4 occupancy in the genomic regions co-occupied by BRD4 and NEUROD1 (Fig. S4, density plot 3 vs. 2), but it did not affect BRD4 occupancy in the genome singly occupied by BRD4 (Fig. S4, density plot 6 vs. 5). Together these results demonstrate that NEUROD1 defines the landscapes of BET bromodomain proteins in SCLC-N cell lines.

### Targeting BET bromodomain proteins leads to suppression of NEUROD1-target genes

We posited that BET bromodomain proteins function as the transcriptional coactivators of NEUROD1 and that inhibiting the former proteins would suppress NEUROD1-target genes. To assess the transcriptional activity of NEUROD1, we first identified its target genes in SCLC. We profiled the transcriptome of three clones of H446 KO and control cells. Through an integrated analysis of the gene expression data and the ChIP-seq data using the Binding and Expression Target Analysis (BETA) algorithm (23), we identified 1,216 NEUROD1-target genes and found that NEUROD1 is primarily a transcriptional activator (Fig. 3A-B). Consistent with a previous report that NEUROD1 plays a crucial role in neuron differentiation (11), Ingenuity pathway analysis identified multiple neuronal-related pathways enriched in the NEUROD1-target genes (Fig. S5).

We next selected a subset of NEUROD1-target genes with at least 4-fold alterations in gene expression upon NEUROD1 KO to constitute a NEUROD1 gene signature (n=384; Fig. 3A). Using the transcriptome data of the SCLC cell lines in the CCLE dataset, we found a significant enrichment of this signature in the SCLC-N lines versus all others (Fig. S6). JQ1 significantly depleted this NEUROD1 gene signature in H446 and COR-L279 cells (Fig. 3C), supporting our hypothesis that the transcriptional activity of NEUROD1 could be blocked by targeting its coactivators - BET bromodomain proteins.

### BETi preferentially suppresses the NEUROD1-target genes associated with superenhancers

To determine what subset of NEUROD1-target genes is more susceptible to BETi, we assessed the effect of JQ1 on the genes with superenhancers (SEs). We compared the BRD4- or NEUROD1- loaded SEs in H446 control cells to the BRD4-loaded SEs in the KO cells (Figs. S7A-C and 3D).

Consistent with our findings that NEUROD1 defines the landscape of BET bromodomain proteins in SCLC, we found in H446 control cells a substantial overlap between the BRD4- and NEUROD1-loaded SEs (Fig. 3D), as well as a strong correlation between the NEUROD1 and BRD4 signals in these SE regions (Fig. S7D). In contrast, only a few SEs are in common between the KO and control cells (Fig. 3D).

Among the NEUROD1-target genes, KO of NEUROD1 caused a more substantial suppression of the genes with NEUROD1-loaded SEs than those without (Fig. 3E), suggesting that these SEs are critical in the regulation of gene expression. Similarly, JQ1 suppressed the genes with the NEUROD1-loaded SEs to a greater degree than those without in H446 control cells (Fig. 3F, left two). In contrast, neither of these gene sets was affected in the NEUROD1-KO cells (Fig. 3F, right two), suggesting these genes became less dependent on the BET family proteins after NEUROD1 KO. To assess whether BET bromodomain proteins became insensitive to JQ1 in the NEUROD1-KO cells, we identified the genes associated with BRD4-loaded SEs in the H446 control (n=307) and KO cells (n=258). These two lists of genes are vastly different but were equally suppressed by BETi, indicating that BET bromodomain proteins were blocked in both H446 KO and control cells (Fig. 3G).

To determine whether BETi suppresses NEUROD1’s transcriptional activity by disrupting the interaction between BET bromodomain proteins and NEUROD1, we performed a co-IP assay after treating H446 cells with JQ1. As shown in Figure 3H, the physical interaction between NEUROD1 and BRD4 remained intact despite JQ1 treatment, suggesting that the interaction between these two proteins is independent of the BD domains. Collectively, these results demonstrate that BETi preferentially suppresses the NEUROD1-target genes associated with SEs.

### The SCLC-N cell lines are more susceptible to BETi

On the basis of the above findings, we predicted that the SCLC-N tumors are more sensitive to BETi. To prove this, we measured *NEUROD1* expression and JQ1 sensitivity in 52 SCLC lines. We found that *NEUROD1* is only expressed in the cell lines sensitive to JQ1 (the cutoff of IC50 to JQ1 = 5µM; Fig. 4A). By grouping all 52 SCLC lines according to their molecular subtypes, we found that the SCLC-N lines were more sensitive to BETi than the SCLC-A lines (Fig. 4B). The differences between the SCLC-N lines and the SCLC-P or SCLC-Y lines had no statistical significance, likely because of too few cell lines in the latter two groups. Using a public high-throughput drug screen dataset (24), we confirmed that the SCLC-N lines are more sensitive to JQ1 than the SCLC-A lines (Fig. S8).

**Figure 4.**
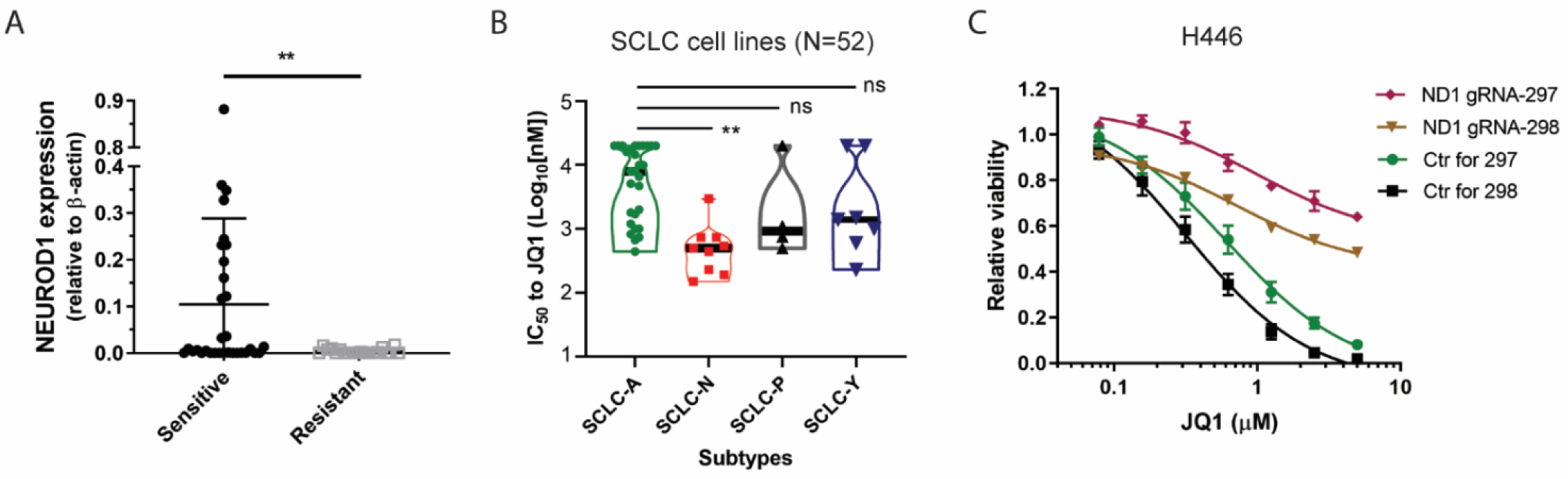
The SCLC-N lines are more susceptible to BETi. **A)** Comparing *NEUROD1* expression between the JQ1-sensitive and -resistant SCLC lines (N=52). An IC50 = 5 µM JQ1 was used as the cutoff to define sensitive and resistant SCLC lines. **B)** Comparing JQ1 sensitivity (IC50) among four molecular subtypes of SCLC lines included in this study (N=52). **C)** Comparing viability between the H446 NEUROD1-KO and control cells following JQ1 treatment (72 hrs). The significance of the two-group comparisons was determined using the Student’s *t*-test (A) and the ANOVA test with Dunnett’s multiple test correction (B). **, P<0.01; ns: not significant.

To further assess the role of NEUROD1 in BETi sensitivity, we compared the response to JQ1 treatment between H446 KO and control cells. As shown in Figure 4C, KO of NEUROD1 led to a more than 20-fold increase in the JQ1’s IC50 in H446 cells. To validate these findings, we knocked out NEUROD1 in DMS-273 (an SCLC-N line) and observed a similar increase in the JQ1’s IC50 value (2.5 µM in the KO cells versus 0.15 µM in control and parental cells; Fig. S9A- B).

MYC expression is known to predict BETi sensitivity in some but not all tumor types. To examine whether NEUROD1 regulates BETi sensitivity through MYC, we first assessed MYC expression after NEUROD1 KO. As shown in Figure S10A-B, KO of NEUROD1 modestly suppressed MYC gene and protein expression in H446 cells but not in DMS-273 cells. Next, we utilized a reporter to assess the effect of BETi on MYC’s transcriptional activity. As shown in Figure S10C, JQ1 suppressed the reporter activity in a dose- and time-dependent manner with no significant distinction between H446 KO and control cells, suggesting that NEUROD1 regulates BETi sensitivity independent of MYC. Collectively, these findings provide supporting evidence that the SCLC-N lines are more susceptible to BETi because of the dependence of NEUROD1 on BET bromodomain proteins for transcriptional activation.

### LSAMP is one of the NEUROD1-target genes mediating BETi sensitivity

To identify the NEUROD1-target gene(s) mediating BETi sensitivity, we reasoned that such gene(s) should also be regulated by BET bromodomain proteins. Therefore, we focused our search among the NEUROD1-target genes affected by BETi (> 2-fold change in expression) and selected 45 genes with a substantial loss of BET bromodomain protein occupancy upon NEUROD1 KO (Fig. 5A). Through siRNA screens, we found 13 and 14 genes consistently showed ≥ 20% growth inhibition after transfection with two distinct siRNAs in H446 and COR-L279 cells (Fig. 5B). Among the ten common candidate genes between these two cell lines (highlighted in red in Fig. 5B), we chose *LSAMP* (limbic system-associated membrane protein) for further investigation because a higher expression of this gene was associated with a worse prognosis in SCLC patients (Fig. 5C). We confirmed that both NEUROD1 and BET bromodomain proteins regulate *LSAMP* gene transcription. As shown in Figure S11A-C, *LSAMP* gene and protein expression decreased substantially in H446 and DMS-273 cells upon NEUROD1 KO. Furthermore, JQ1 decreased LSAMP protein expression in a dose-dependent manner in H446 cells (Fig. S11D).

**Figure 5.**
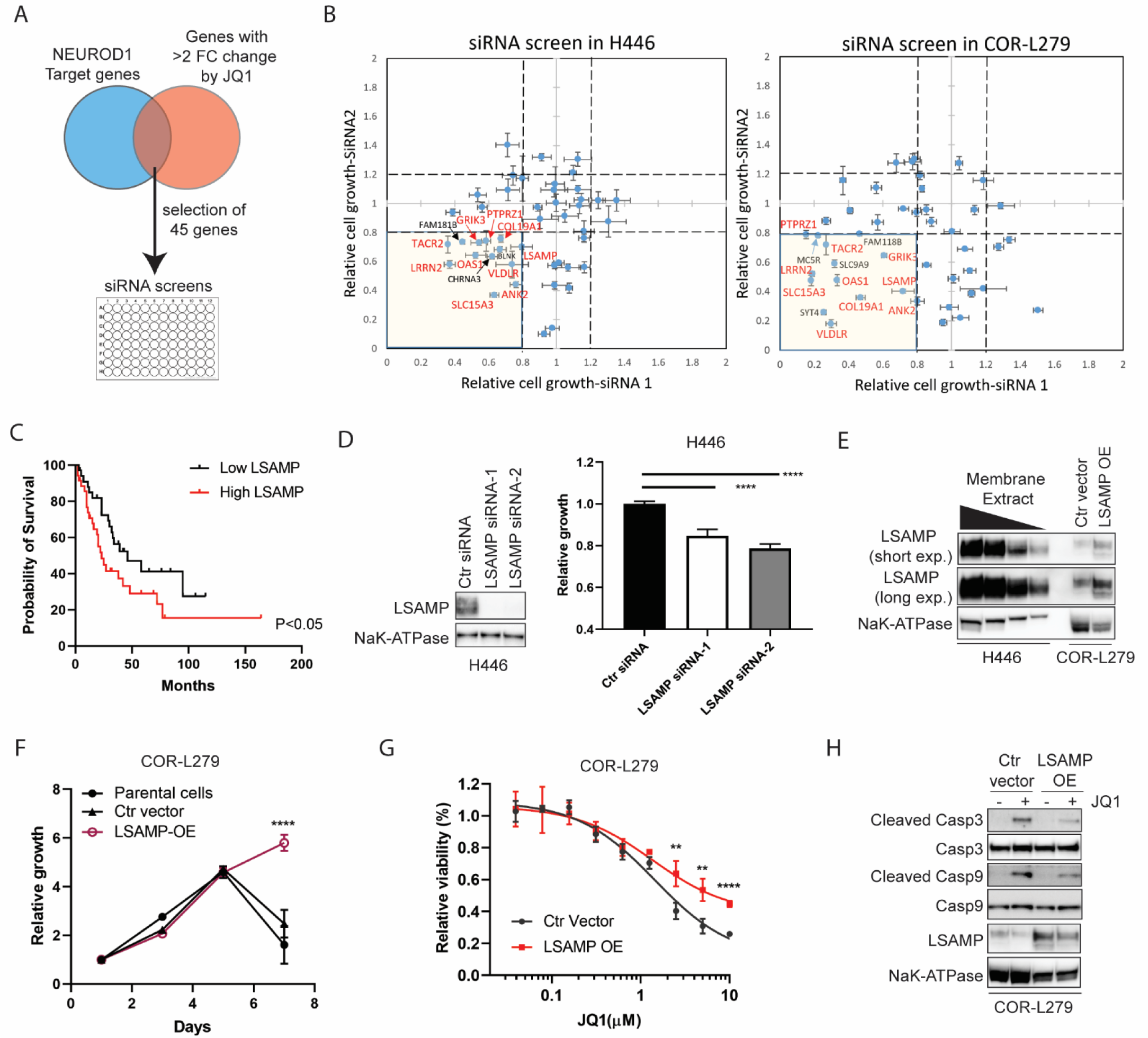
LSAMP, a NEUROD1-target gene, partially mediates BETi sensitivity. **A)** A diagram showing the selection of 45 genes regulated by both NEUROD1 and BET bromodomain proteins for siRNA screens. **B)** The results of siRNA screens (72 hrs) in H446 (left) and COR- L279 cells (right). The genes highlighted in red are the common genes between H446 and COR- L279 cells that consistently showed ≥ 20% growth inhibition upon knockdown with two distinct siRNAs. **C)** Kaplan-Meier curves showing an inverse correlation between *LSAMP* expression and the overall survival in SCLC patients (N=69; ref. (38)). The patients were dichotomized on the basis of *LSAMP* transcript levels in the tumors (top 50% vs. bottom 50%). **D)** LSAMP knockdown suppresses H446 proliferation. LSAMP was knocked down using two siRNAs distinct from those used in the initial screens. Left, LSAMP expression in the membrane protein fractions. NaK- ATPase serves as a loading control. Right, the effect of LSAMP knockdown on H446 cell growth, as measured by the CellTiter Glo 2.0 assay 72 hrs after transfection. **E)** Stable expression of ectopic LSAMP in COR-L279 cells. A tapering amount of membrane protein from H446 (20, 10, 5, 2.5 µg) was loaded to compare with 20 µg membrane protein from COR-L279 cells. **F)** Comparing the proliferation rate of the COR-L279 cells with ectopic LSAMP to control or parental cells. **G)** Differential sensitivity to JQ1 (72 hrs) between the COR-L279 with ectopic LSAMP and control cells. **H)** Western blots showing decreased caspase 3 and 9 cleavages after JQ1 treatment (5µM, 24 hrs) in the COR-L279 with ectopic LSAMP compared to control cells. The significance of two-group comparisons was determined using the Log-rank test (C), the ANOVA test with Dunnett’s multiple test correction (D, F), and the Student’s *t*-test (G). **, P<0.01; ****, P<0.0001. Ctr, control; OE, overexpression.

To assess its function in SCLC, we knocked down *LSAMP* using two siRNAs distinct from the ones used in the screens (Fig. 5D left panel) and found a similar degree of growth inhibition as in the initial screens (Fig. 5D right panel). Next, we stably overexpressed LSAMP in COR-L279, which has a much lower expression of this gene compared to H446 (Fig. 5E). Although the COR- L279 cells with ectopic *LSAMP* did not grow faster than control cells, these cells continued to proliferate after day 5, while the control and parental cells showed decreased viability (Fig. 5F). Furthermore, the COR-L279 cells with ectopic LSAMP were less susceptible to high doses of BETi than control cells (Fig. 5G). On the basis of these results, we posited that ectopic expression of LSAMP rendered cells less prone to apoptosis. Consistent with this notion, JQ1 caused less cleavage of caspase 3 and caspase 9 in the cells with ectopic LSAMP compared to control (Fig. 5H). Together these results demonstrate that *LSAMP* is one of the NEUROD1-target genes mediating BETi sensitivity.

### BETi inhibits the NEUROD1 gene network and suppresses SCLC growth in vivo

To assess the therapeutic effects of BETi *in vivo*, we chose NHWD-870 (a potent BETi in early phase clinical trials) over JQ1 because the latter has unfavorable pharmacokinetic properties *in vivo* (25). We first confirmed that NHWD-870 has a similar sensitivity profile as JQ1 among SCLC lines (Fig. S12). Using an SCLC-N PDX model - LX22, we found NHWD-870 significantly decreased tumor burden and extended median survival of the tumor-bearing mice from 27 to 42 days (Fig. 6A). To assess how NHWD-870 affected NEUROD1-target genes in LX22 tumors, we performed RNA-seq to profile the transcriptome changes in the tumor cells isolated from the xenografts following one-week NHWD-870 treatment. As shown in Figure 6B-C, NHWD-870 significantly depleted the NEUROD1 gene signature and decreased *LSAMP* transcript levels in the xenograft tumors without affecting *NEUROD1* expression.

**Figure 6.**
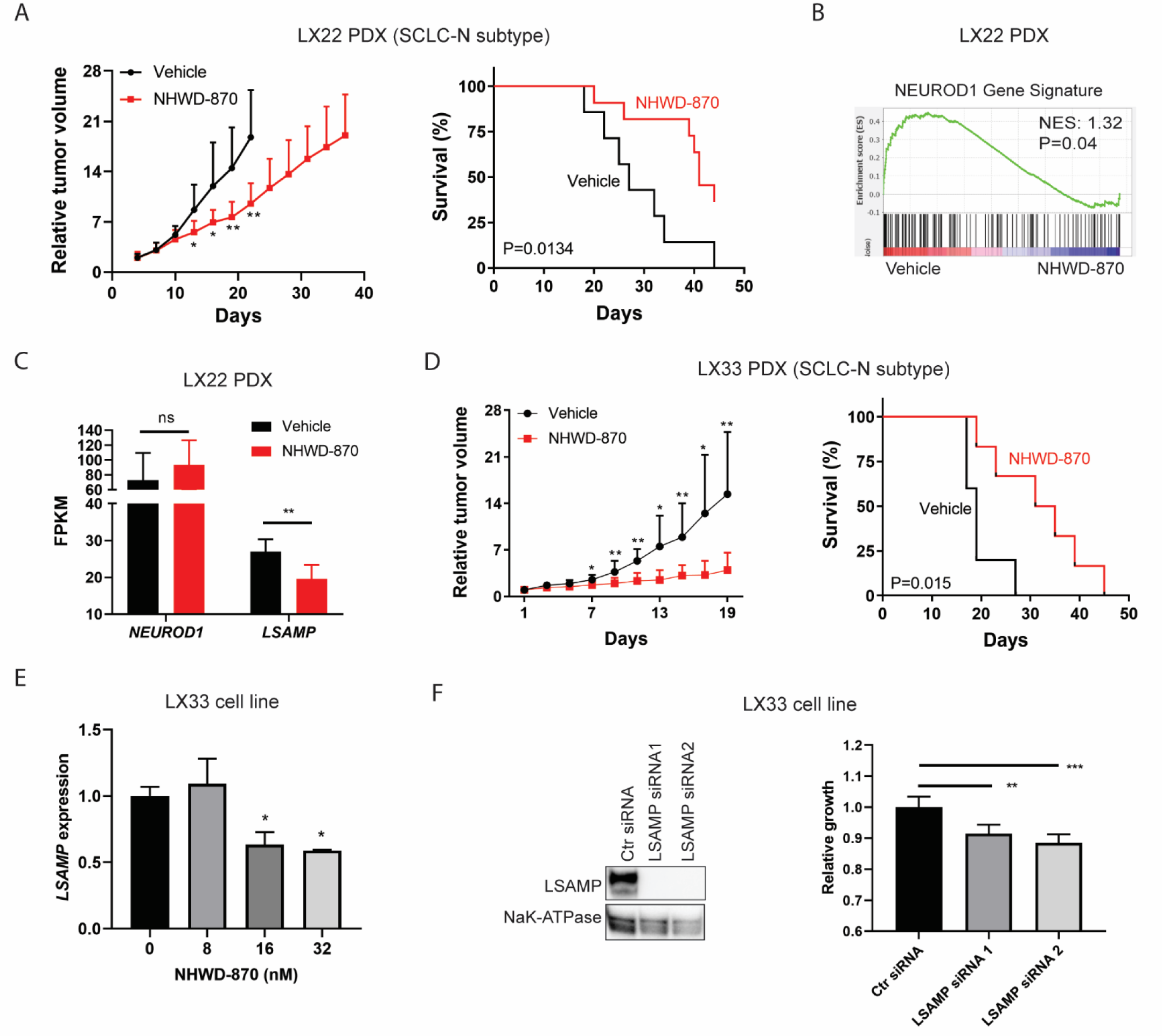
BETi suppresses NEUROD1 transactivation and inhibits SCLC growth *in vivo*. **A)** NHWD-870 (2mg/kg via gavage on a 5-day-on/2-day-off schedule) suppressed the tumor growth (left) and extended the survival (right) in the LX22 SCLC PDX model (an SCLC-N subtype). n=13 per group. **B)** A GSEA plot showing that NHWD-870 (2mg/kg via gavage daily for five days) significantly depleted the NEUROD1 gene signature in the LX22 PDX xenografts. n=6 per group. **C)** Comparing *NEUROD1 and LSAMP* transcript abundance (FPKM) in xenografts treated with NHWD-870 versus the vehicle in the LX22 model. **D)** NHWD-870 (3mg/kg via gavage daily) suppressed the growth (left) and extended the survival (right) in the LX33 SCLC PDX model (an SCLC-N subtype). n=6 per group. **E)** NHWD-870 (24 hrs) caused a dose-dependent decrease of *LSAMP* expression in LX33 cells. Gene expression was measured by qRT-PCR. **F)** LSAMP is essential for LX33 growth. Left, knockdown of LSAMP by two distinct siRNAs. Right, the effects of LSAMP knockdown on the growth of LX33 cells (72 hours post-transfection). The significance of the two-group comparison was determined using the Student’s *t*-test (A and D, left panels; C), the Log-rank test (A and D, right panels), and the ANOVA test with Dunnett’s multiple test correction (E, and F right panel). *, P<0.05; **, P<0.01; ***, P<0.001; ns, not significant. Ctr, control; FPKM, Fragments Per Kilobase of transcript per Million mapped reads; NES, normalized enrichment score; PDX, patient-derived xenograft.

To confirm these findings, we utilized another SCLC-N PDX model - LX33. Like LX22, NHWD-870 significantly lessened the tumor burden and extended the median survival in this model (Fig. 6D). We attempted to establish tumor cell lines from these two PDX models but could only succeed with LX33. In the LX33 SCLC line, NHWD-870 suppressed *LSAMP* expression as low as 16nM (Fig. 6E). LSAMP knockdown decreased LX33 cell proliferation, supporting that BETi inhibits SCLC growth by suppressing the NEUROD1 gene network *in vivo* (Fig. 6F).

## Discussion

Because the master TFs define the transcriptome landscapes of SCLC, it is appealing to target these genes for therapeutic purposes. Here, we present evidence that BET bromodomain proteins partner with NEUROD1 to activate the transcription of its downstream genes, resulting in the susceptibility of the SCLC-N tumors to BET inhibitors. Our results demonstrate that targeting transcriptional coactivators could be a new approach to block the functions of the master TFs and has the potential to be developed into a subtype-specific therapy.

Consistent with the role of NEUROD1 as a master TF in SCLC, our results demonstrated that it physically interacts with BET bromodomain proteins and defines the genomic landscapes of the latter proteins in H446 cells. The BETi resistance in the NEUROD1-KO cells is likely because of the altered landscapes of BET bromodomain proteins, causing these factors to be associated with a largely distinct set of genes. Although BETi was equally effective in inhibiting BET bromodomain proteins, the resistance of the NEUROD1-KO cells to BETi suggests that these genes are not essential for cell growth.

In search of the genes mediating BETi sensitivity in SCLC-N tumors, we identified *LSAMP* as one of the potential candidates through the siRNA screens. *LSAMP*, a member of the IgLON family of immunoglobulin domain-containing cell adhesion molecules, is considered a tumor suppressor gene in ovarian cancer and osteosarcoma on the basis of its frequent deletion and a positive correlation between *LSAMP* expression and the overall survival in these malignancies (26, 27). However, in SCLC, a higher expression of this gene was correlated with a worse prognosis, and ectopic expression of this gene decreased apoptosis. These findings suggest that the functions of LSAMP could be tumor type-specific. Besides *LSAMP*, our siRNA screen results indicate that many other NEUROD1-target genes suppressed by BETi also play a role in tumor growth. Therefore, BETi inhibits SCLC growth likely by suppressing a panel of NEUROD1-target genes. In contrast to a previous report that ASCL1 expression could predict BETi sensitivity in SCLC (28), the results from this study and a high-throughput screen dataset showed the SCLC-A lines are generally more resistant to BETi than the SCLC-N lines. Although BETi decreases ASCL1 expression (28), this could lead to activation of IGF1 signaling due to the suppression of IGFBP5, a negative regulator of IGF1 signaling and an ASCL1-target gene (29). This feedback mechanism was not present in SCLC-N lines (29), which may explain the primary resistance of SCLC-A lines to BETi.

It has been recently recognized that SCLC has substantial intratumoral heterogeneity, which could be a potential obstacle for subtype-specific therapies. Two recent studies that assessed molecular subtype composition in SCLC tumors by immunohistochemical staining found that 20.4-37% of the tumors had two or more subtypes (6, 7). Since our results showed that the non-SCLC-N subtypes are less sensitive to BETi, BETi single-therapy may select for these subtypes and result in early recurrence. One potential solution to this obstacle is to use drug combinations that would be effective against two or more subtypes. For instance, in another study, we have found that mTOR inhibitors could potentiate the antitumor activities of BETi and make the latter drug effective in the BETi-resistant SCLC lines (30).

In summary, this study demonstrates that SCLC-N subtype tumors are susceptible to BETi due to the dependence of NEUROD1 on BET bromodomain proteins for transcription activation. Our results suggest that blocking transcriptional coactivator(s) is an alternative approach to targeting the master TFs in SCLC and has the potential to be developed into subtype-specific therapies.

## Materials and Methods

### Cell Culture

All SCLC lines used in this study were purchased from commercial vendors (Table S1). Cells were cultured in recommended media and maintained in a humidified incubator at 37 °C with 5% CO2. LX33 SCLC line was established by culturing tumor cells isolated from an LX33 SCLC PDX in HITES media (media composition is listed in Table S1). All cell lines tested negative for mycoplasma. The short tandem repeat analysis was performed to authenticate commercial cell lines.

### CRISPR Knockout

CRISPR was performed using a standard two-step method (31). Briefly, a Cas9 expression vector, lentiCas9-Blast (Addgene, #52962; a gift from Feng Zhang (32)), was transfected into H446 and DMS-273 cells by lipofection. Cells were selected with 1-2.5 µg/ml blasticidin (Gibco). Subsequently, the Cas9-expressing cells were infected with lentiviral particles containing gRNA-carrying vectors targeting NEUROD1 (Sigma): HS5000024297 (short as #297), GAGTCCTCCTCTGCGTTCATGG; HS5000024298 (#298), GAGGAGGAGGACGAAGATGAGG. Transduction was performed at a multiplicity of infection (MOI) of 2 in the presence of 5 µg/ml polybrene (Sigma). Cells were selected with 0.5-2.5 and µg/ml puromycin (Gibco), and KO of NEUROD1 was verified by western blot and Sanger sequencing. The NEUROD1-KO cells were maintained in complete media without selection antibiotics after verification.

### RNA Sequencing (RNA-seq)

Total RNA libraries (COR-L279) and mRNA libraries (H446 and LX22) were constructed using a TruSeq Stranded Total RNA Kit (Illumina, RS-122-2201) and an mRNA Library Prep kit V2 (Illumina, RS-122-2001), respectively. Paired-end sequencing of the constructed libraries was performed on a HiSeq 4000 system (Illumina), and the raw fastq data were analyzed using the CCBR/Pipeliner (33). A false-discovery rate of 0.05 was set as a cutoff to define significant alterations in gene expression.

### Gene Set Enrichment Analysis (GSEA)

GSEA was performed using GSEA (v.4.0.2) under the setting of gene_set permutation at 1,000 times, with the DESeq2 normalized counts as data input. The NEUROD1 gene signature comprises 384 NEUROD1-target genes that showed at least 4-fold alterations in gene expression upon NEUROD1 KO in H446 cells (Table S2).

### ChIP-sequencing (ChIP-seq)

ChIP-seq was performed as described previously (34). For each assay, 4 − 10 µg of the following antibodies were used: α-NEUROD1, Cell Signaling Technology, #4373; α-BRD2, Cell Signaling Technology, #5848; α-BRD3, Bethyl, A302-368A; α-BRD4, Bethyl, A700-004; α-H3K27Ac, Abcam, #ab4729. The procedures of ChIP-seq analysis were provided in supplemental methods.

### siRNA Transfection and Screen

LSAMP siRNAs (#S8299 and #S8301; ThermoFisher Scientific) were transfected into cells using Lipofectamine RNAiMax (Thermo Scientific) at a final concentration of 20nM. The siRNA library that contained 90 siRNA targeting 45 genes of interest was custom synthesized by Qiagen (Table S3).

### Plasmid Transfection and Establishment of Stable Transformant

Plasmids were transfected into cells using Lipofectamine 3000 reagent (ThermoFisher Scientific) following a standard protocol. LSAMP stable transformants were established by transfecting COR-L279 cells with a tagged ORF clone of this gene (Origene; # RC207618) followed by 1mg/ml G418 selection (Roche) in semi-solid media (ClonaCell™-TCS medium, Stem Cell Technology). The surviving cell clones were expanded and maintained in complete media with 0.4 mg/ml G418.

### Animal Studies

The animal experiments were approved and performed according to the regulations set by the National Cancer Institute-Bethesda Animal Care and Use Committee. SCLC PDX LX22 and LX33 models were generous gifts from John T. Poirier and Charles M. Rudin (35, 36). Freshly isolated PDX tumors were dissociated into single-cell suspension using a Human Tumor Dissociation Kit (Miltenyi Biotec) and a gentleMACS™ Octo Dissociator (Miltenyi Biotec). After lyzing red blood cells with ACK lysis buffer (Quality Biological), the remaining cells were washed with ample PBS three times and resuspended in PBS at 5x10^7^ viable cells per ml. One hundred µl of cell suspension (∼ 5x10^6^ viable cells) was injected subcutaneously into right flanks of 6-week-old NOD-SCID mice (Charles River Laboratories). Once the tumor volume reached between 50 and 100 mm^3^ (average, 14-21 days), mice were randomized, and drug treatments were given at the doses and frequencies specified in the figure legends. Animals were euthanized if 1) tumor volume was ≥1500 mm^3^; 2) tumor became ulcerated; 3) 75 days had elapsed after a tumor became palpable. For the transcriptome analysis, tumor cells were separated from mouse stromal cells using a magnetic-activated cell sorting method (37), and RNA was extracted using an RNeasy Mini Kit (Qiagen).

For tumorigenicity assay, H446 NEUROD1-KO cells (gRNA-297, cl.12 and cl.15) and control cells (cl.10) were resuspended in Matrigel (Corning). 1x10^6^ cells in 100 µl were inoculated subcutaneously into each flank of athymic mice. Tumors were resected at three weeks (control and NEUROD1 gRNA-297 cl.15) or eight weeks (NEUROD1 gRNA-297 cl.12) post-injection.

## Acknowledgments

This study was supported by the Center for Cancer Research, the Intramural Program of the National Cancer Institute (NCI) (grant ZIA BC011787: Chen), and in part by an NCI FLEX award (grant ZIA BC011839: Chen and Schrump) and NCI U01 CA231844 (Oliver). The SCLC PDX models were made available through the support of NCI U24 CA213274 (Charles Rudin). The authors thank Yulong Li, Yan-Jin Liu, Shih-Hsin Hsiao, Kaitlin C. McLoughlin, and Sichuan Xi for their assistance in this study; and Nenghui Wang for providing NHWD-870.

## Supplemental Methods

### RNA Isolation and Quantitative real-time PCR Assay

RNA was extracted from cells using an RNeasy Mini Kit (Qiagen), and cDNA was synthesized using an iScript cDNA Synthesis Kit (Bio-Rad). Quantitative gene expression analysis was performed on an ABI PRISM7500 real-time PCR system (ThermoFisher), using either TaqMan Fast Advanced Master Mix or SsoAdvanced™ Universal SYBR® Green Supermix (Biorad). The predesigned specific primers were purchased from ThermoFisher Scientific, Biorad, and Integrated DNA Technologies.

### ChIP-seq – Peak Calling

An initial quality check of the fastq files was performed using FastQC, and adapter sequences and low-quality base pairs were trimmed using Cutadapt. Trimmed reads were mapped to the human hg19 reference genome using the backtrack algorithm of Burrows-Wheeler Aligner (BWA-Backtrack) (1). Duplicated reads were filtered, and the peaks were called using Model-based Analysis of ChIP-Seq (MACS) with the corresponding input as control (2). Peaks of BRD2, BRD3, and BRD4 were called using default parameters, and NEUROD1 peak calling was performed using the parameters optimized for transcription factors (2). Genomic annotations of the NEUROD1 peaks were performed using the ‘annotatePeaks’ function of the Homer v4.11 (3).

### ChIP-seq – Metagene Analysis

The metagene profile of the NEUROD1 peaks was performed using the ‘makeMetaGeneProfile’ function of the Homer, using the NEUROD1 peaks as the input.

### ChIP-seq – Peak Colocalization

Tag directories were first generated from the bam files of ChIP-seq samples and the corresponding input, using the ‘makeTagDirectory’ function of the Homer. Histograms on BET bromodomain protein occupancy within 2.5 KB bilateral of the centers of NEUROD1 peaks were generated using the Homer’s ‘annotatePeaks’ function with the parameter ‘-size 5000 -hist 10’. The results were plotted using GraphPad Prism (v8.4.3).

### ChIP-seq – Heatmap

BRD4 peaks colocalizing with NEUROD1 peaks were identified using Homer’s ‘mergePeaks’ function with the setting ‘-d given’. Peak annotation was performed using Homer’s ‘annotatePeaks’ function. The NEUROD1 or BRD4 occupancy near the BRD4 peaks was assessed using Homer’s ‘annotatePeaks’ function, and the results were visualized in heatmaps using EaSeq (4).

### ChIP-seq – Superenhancer (SE) Analysis

The H3K27Ac peaks were used to mark constituent enhancer sites in the H446 control cells and the NEUROD1-KO cells, and BRD4-loaded or NEUROD1-loaded SEs were called by ROSE using the default setting (5, 6). The identified SEs were annotated for the nearest genes using Homer’s ‘annotatePeaks’ function.

### ChIP-seq – de novo Motif Discovery

Motif discovery was performed using the ‘findMotifsGenome’ function of Homer v4.11, with the setting of ‘-size 200 -mask’ and the NEUROD1 peaks as the input.

### Identification of NEUROD1-target Genes

The NEUROD1-target genes were identified using the Binding and Expression Target Analysis (BETA) algorithm (7). Input data included NEUROD1 peaks from the ChIP-seq analysis and differential gene expression output from Limma (H446 NEUROD1-KO cells vs. control cells). The analysis was performed with the ‘Plus’ option in the setting of ‘--df 0.05 --bl --da 0.5’.

### Western Blot Analysis

Whole-cell lysates were extracted using RIPA lysis buffer (Millipore), and membrane proteins were isolated using a Mem-PER Plus Membrane Protein Kit (ThermoFisher). Western blots were performed using a standard immunoblotting protocol. The antibodies and their dilutions were: NEUROD1 (BD Biosciences, #563000, 1:2,000), BRD2 (Santa Cruz Biotechnology, #sc-393720, 1:200), BRD3 (Santa Cruz Biotechnology, #sc-81202, 1:200), BRD4 (Novus, NBP2-52959, 1:1,000), LSAMP (Abcam, #ab89719, 1:1,000), NaK-ATPase (Santa Cruz Biotechnology, #sc-71638, 1:1,000), cleaved caspase 3 (Cell Signaling Technology, #9661, 1:1,000), caspase 3 (Cell Signaling Technology, #9662, 1:1,000), cleaved caspase 9 (Cell Signaling Technology, #7237, 1:1,000), caspase 9 (Cell Signaling Technology, #9502, 1:1,000), MYC Cell Signaling Technology, # 5605, 1:1,000); γ-tubulin (Sigma, #T6557, 1:10,000).

### Co-immunoprecipitation (Co-IP) Assays

Nuclear extracts were prepared from H446 and H524 cells using a Nuclear Complex Co-IP Kit (Active Motif, #54001). Each IP, in a final volume of 500 µl, contained 200-500 µg nuclear extract, an antibody against the protein of interest (αBRD4, Bethyl, A301-985A, 5 µg; αNEUROD1, Cell Signaling Technology, #4373, 10 µl; αBRD2, Cell Signaling Technology, #5848, 10 µl) or control IgG (Cell Signaling Technology, #3900, 5 µg), 20 µl slurry of Magna ChIP protein A magnetic beads (Millipore), and 1x IP High buffer supplemented with a protease inhibitor cocktail. After overnight mixing on an orbital shaker at 4 °C, antibody/antigen/bead complexes were washed three times with 1x High Buffer supplemented with 1mg/ml bovine serum albumin (BSA), followed by three washes with 1x High Buffer without BSA. The immunoprecipitated protein complexes were eluted by mixing the washed beads with Laemmli buffer supplemented with 5% 2-mercaptoethanol (Sigma) after heating at 95 °C for 5 min. The presence of the protein of interest was detected by western blot.

### Cell Proliferation Assay and Viability Assay

After trypsinization, cell suspensions were passed through 40 µm cell strainers. Five thousand cells in 100 µl growth media were seeded into each well of a 96-well white plate with a clear bottom (Corning, #3610). Alternatively, 750 cells in 15 µl were seeded into each well of a 384-well white plate with a clear base (Corning, #3765).

For cell proliferation assay, cell viability was checked the next day (D1) and then every two days using CellTiter-Glo® 2.0 Reagent (Promega). A reflective foil seal (Bio-Rad, #MSF1001) was applied to the bottom of each plate to maximize the signal. The results were normalized to the reading on day 1. Four replicates were used for each data point.

For cell viability assay, JQ1 (Sigma) or NHWD-870 (a kind gift from Nenghui Wang) were serially diluted 2-fold in DMSO to generate nine consecutive concentrations. The compounds were first diluted 100 times in culture media, and the diluted drug solution was then delivered to the cells at 10% of the final volume to achieve a total of 1,000-time dilutions. After brief mixing, cells were incubated for 72 hours before cell viability was measured. Four replicates were used for each dose of drug treatment.

### MYC reporter assay

An MYC Firefly luciferase reporter (pBV-Luc wt MBS1-4; Addgene plasmid #16564; a gift from Bert Vogelstein (8)) and a Renilla reporter (pRL-CMV; Promega) were transfected into H446 NEUROD1-KO and control cells at 8:1 ratio by lipofection. One day after transfection, the cells were trypsinized and seeded into a 384-well plate at 750 cells per well, and JQ1 treatment was administered subsequently. The Firefly and Renilla luciferase activities were measured at 24- and 48-hour intervals using a Dual-Glo Luciferase assay system (Promega). Each dose had four replicates.

### Statistical Analysis

Statistical analyses and graphing were performed using GraphPad Prism. All tests were performed with a two-sided significance level of 0.05.

### Data availability

All data sets described in the paper are deposited in NCBI Gene Expression Omnibus (accession number: pending).

## Supplemental tables

**Figure S1.**
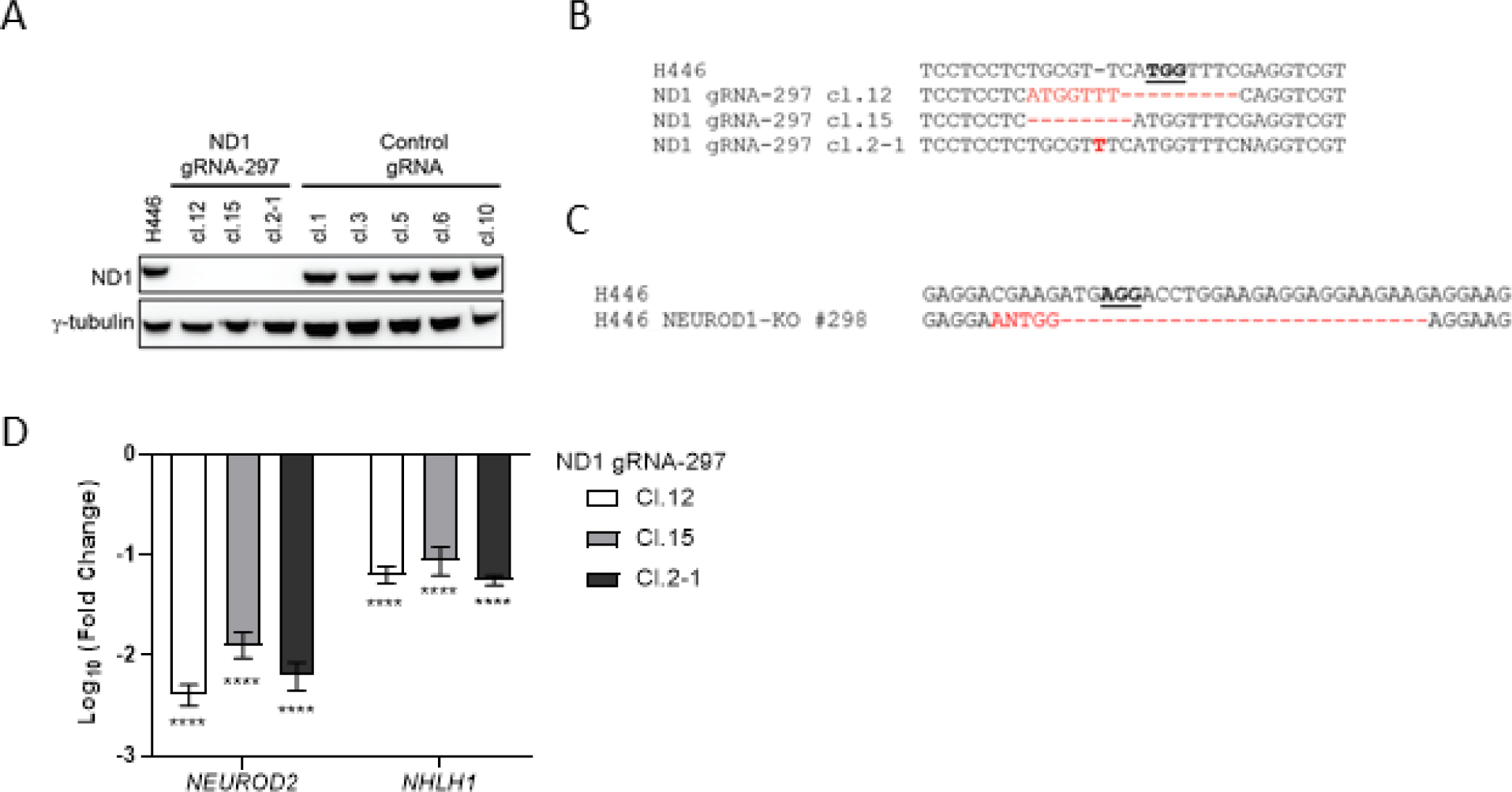
Knockout of NEUROD1 in H446 cells by CRISPR. **A)** Western blots showing a complete loss of NEUROD1 protein expression in three NEUROD1-KO clones (cl. 12, cl. 15, and cl. 2-1) receiving the gRNA #297. **B)** Sanger sequencing identified indels (highlighted in red) in the NEUROD1 genomic DNA of three H446 clones receiving the gRNA #297. The PAM sequence was underlined. **C)** An indel was detected in the NEUROD1 genomic DNA of the H446 clone receiving the gRNA #298. **D)** Expression of two NEUROD1-target genes (NEUROD2 and NHLH1) was measured in the three H446 NEUROD1-KO cells (gRNA-297: cl. 12, cl. 15, and cl. 2-1) using quantitative real-time reverse-transcription PCR.

**Figure S2.**
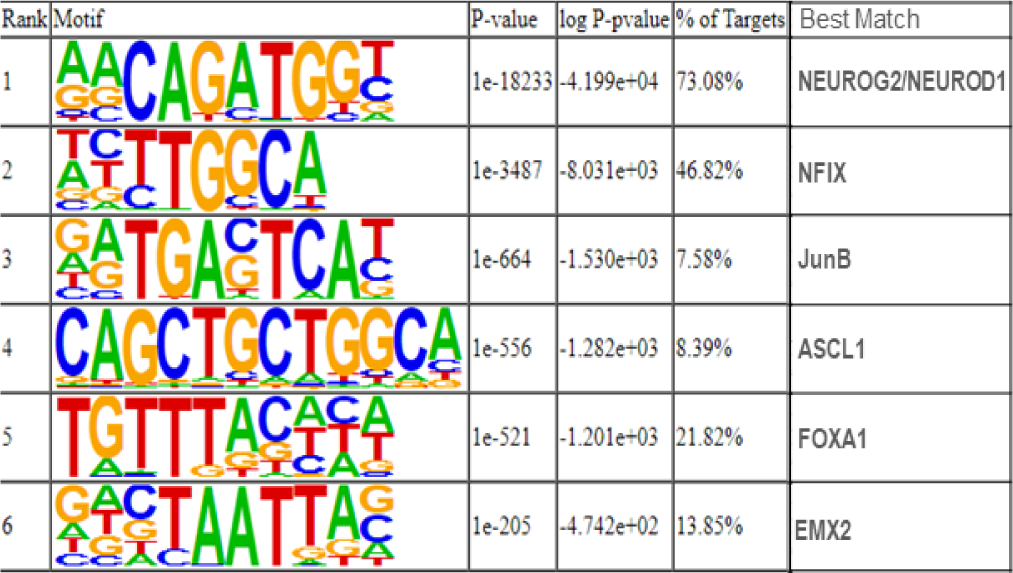
The top six motifs identified in the genomic sequences corresponding to the NEUROD1 peaks in H446 control cells. De novo motif discovery analysis was performed using Homer.

**Figure S3.**
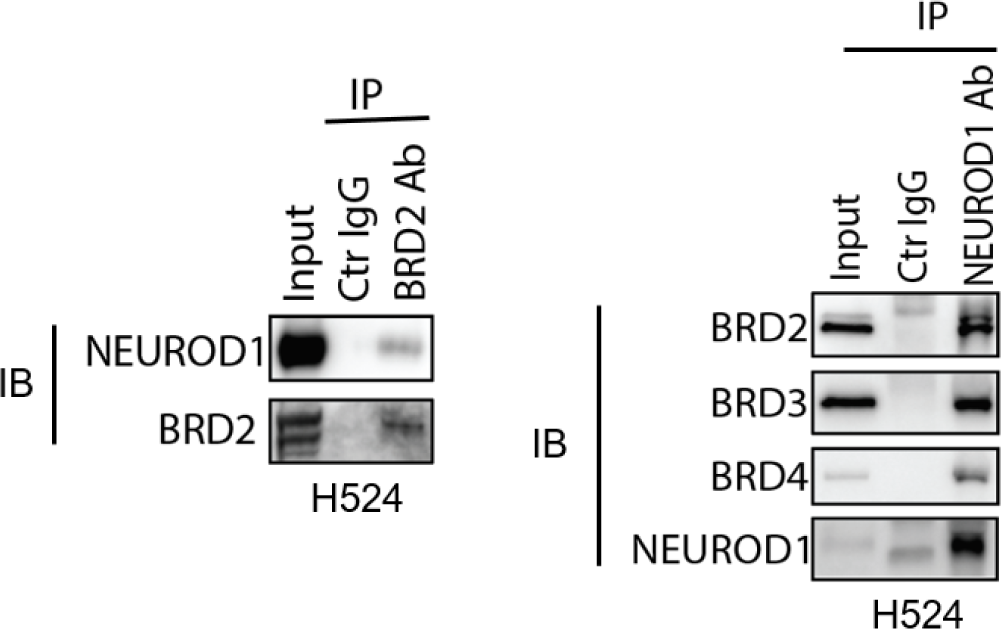
BET bromodomain proteins interact with NEUROD1 in H524 cells. Nuclear extracts isolated from H524 cells were subject to BRD2 (left) or NEUROD1 (right) immunoprecipitation, and the presence of NEUROD1 (left) and BET bromodomain proteins (right) in the elute was detected using western blot. Ab, antibody; IB, immunoblotting; IP, immunoprecipitation.

**Figure S4.**
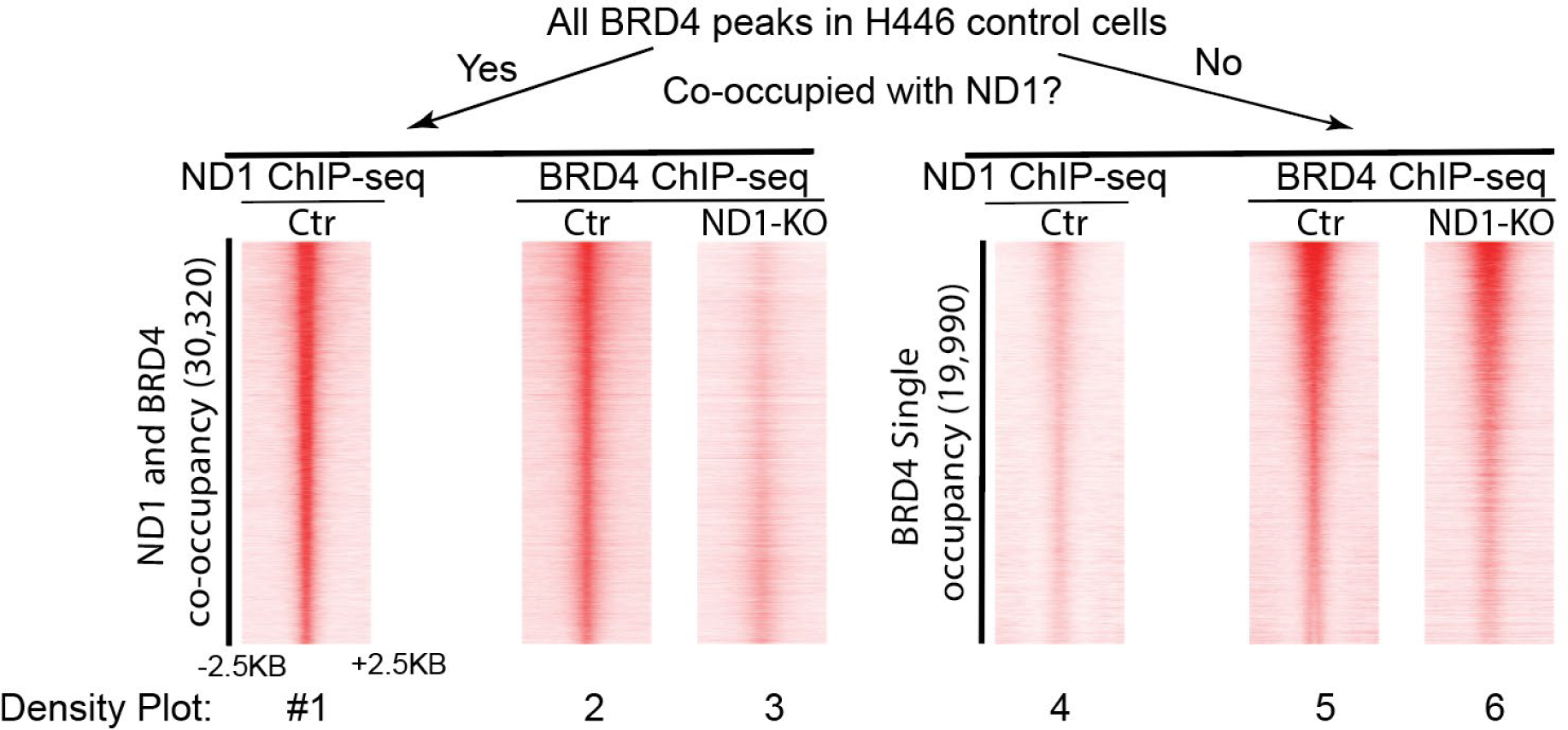
The NEUROD1 KO affects the BRD4 occupancy only in the genomic regions co-occupied by NEUROD1. BRD4 peaks in H446 control cells were separated into two groups on the basis of NEUROD1 co-occupancy (density plot #1 vs. #4). NEUROD1 KO affected BRD4 occupancy in the NEUROD1/BRD4- colocalized genomic regions (#2 vs. #3) but not in the BRD4 singly-occupied regions (#5 vs. #6).

**Figure S5.**
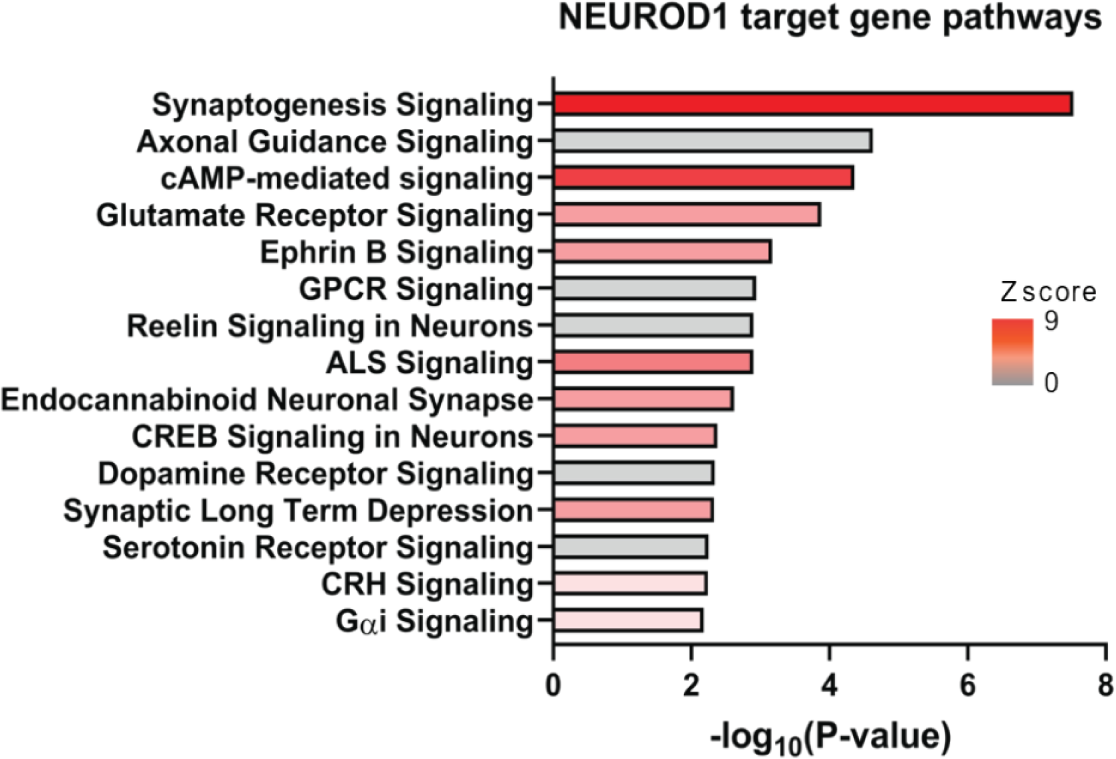
Neuronal-related genes are enriched in the NEUROD1-target genes in H446 cells. Ingenuity Pathway Analysis identified 15 pathways enriched in the NEUROD1-target genes. ALS, amyotrophic lateral sclerosis; CREB, cAMP response element-binding protein; CRH, corticotrophin-releasing hormone; GPCR, G protein-coupled receptor.

**Figure S6.**
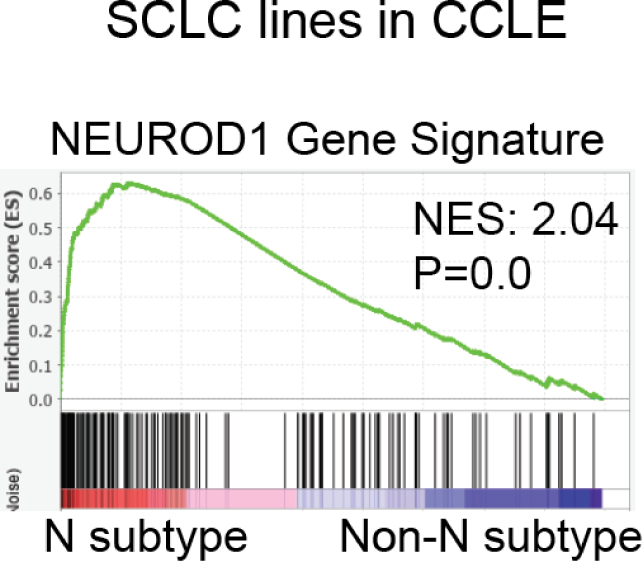
Validation of the NEUROD1 gene signature using the SCLC RNA-seq data from the CCLE dataset. A GSEA plot showing enrichment of the NEUROD1 gene signature in the SCLC-N cell lines (N subtype) compared to all other SCLC lines (non-N subtype) included in the CCLE dataset.

**Figure S7.**
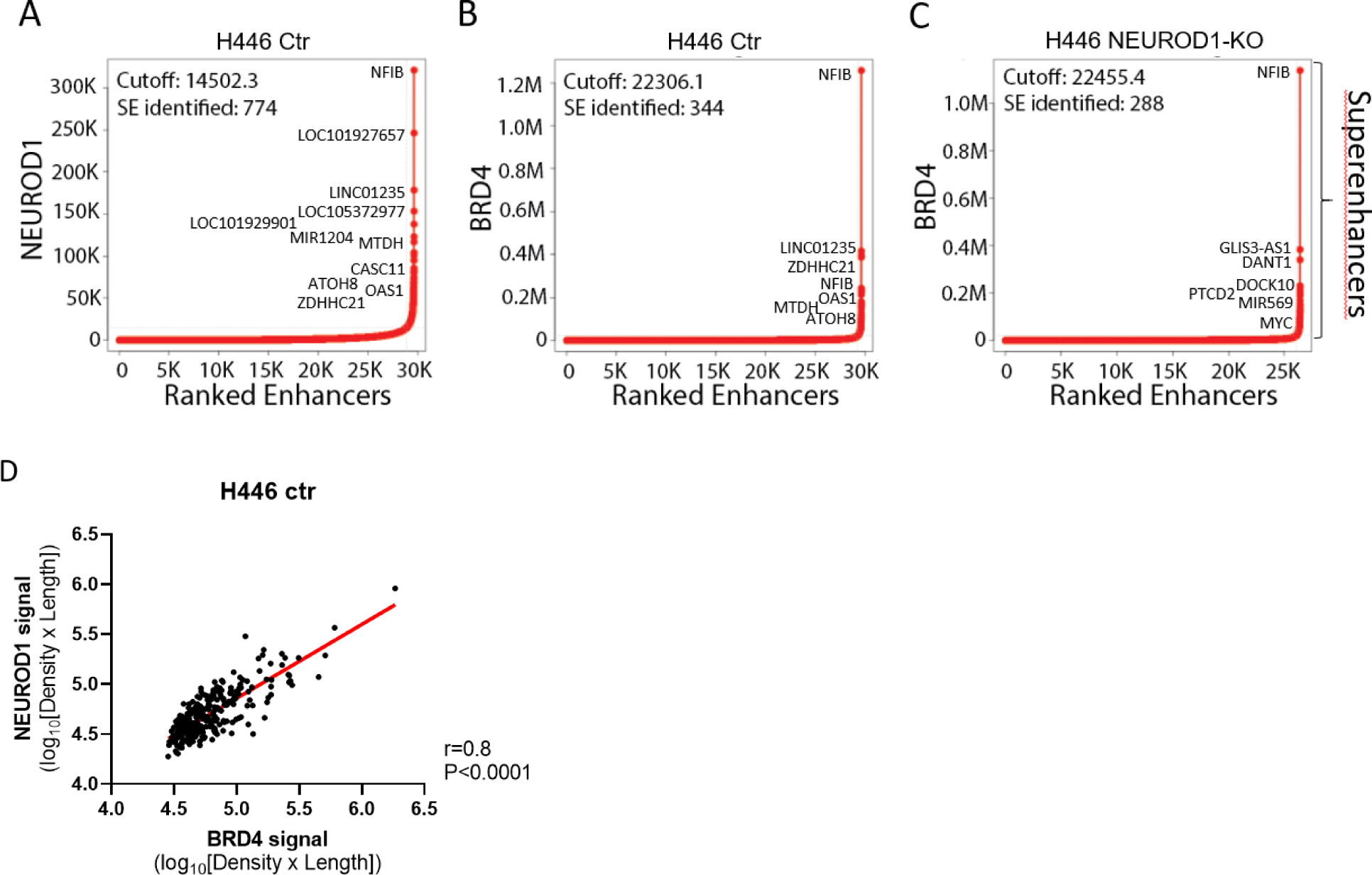
The NEUROD1- and BRD4-loaded SEs identified in H446 control and NEUROD1-KO cells. **A-B)** Stitched NEUROD1 (A) and BRD4 (B) enhancers in increasing rank order based on ChIP-seq signal minus the input signal in H446 control cells. **C)** Stitched BRD4 enhancers in increasing rank order in the H446 NEUROD1-KO cells. **D)** Correlation between NEUROD1 and BRD4 signals (Density x Length) at the SEs co-occupied by these two proteins in the H446 control cells. Pearson correlation coefficient and P value were shown.

**Figure S8.**
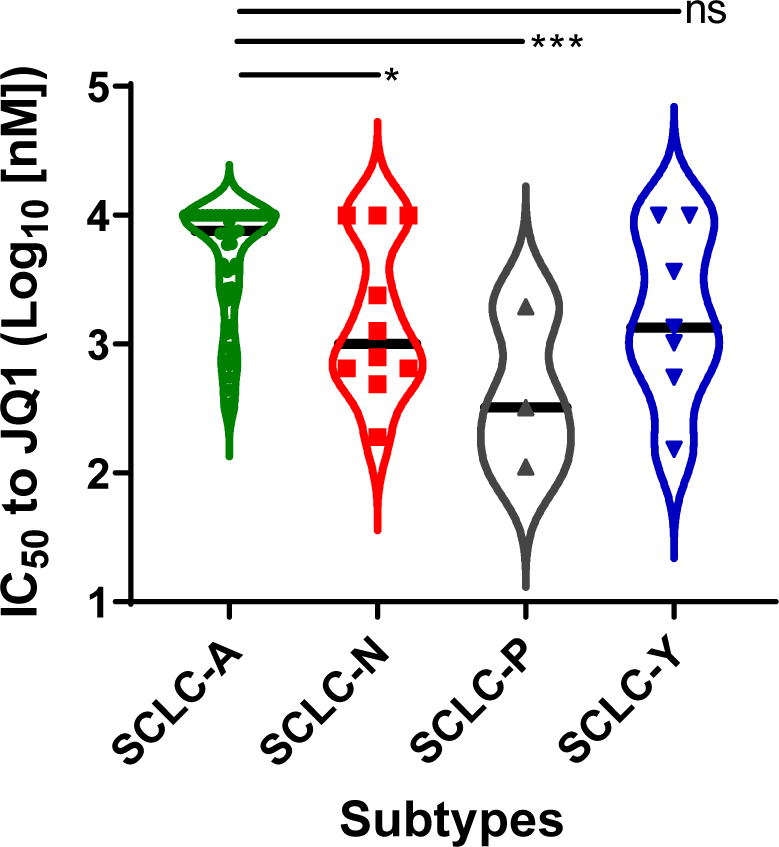
Comparison of JQ1 sensitivity (IC50) among the four molecular subtypes of SCLC cell lines (N=57) in a public high-throughput drug screen dataset (24).

**Figure S9.**
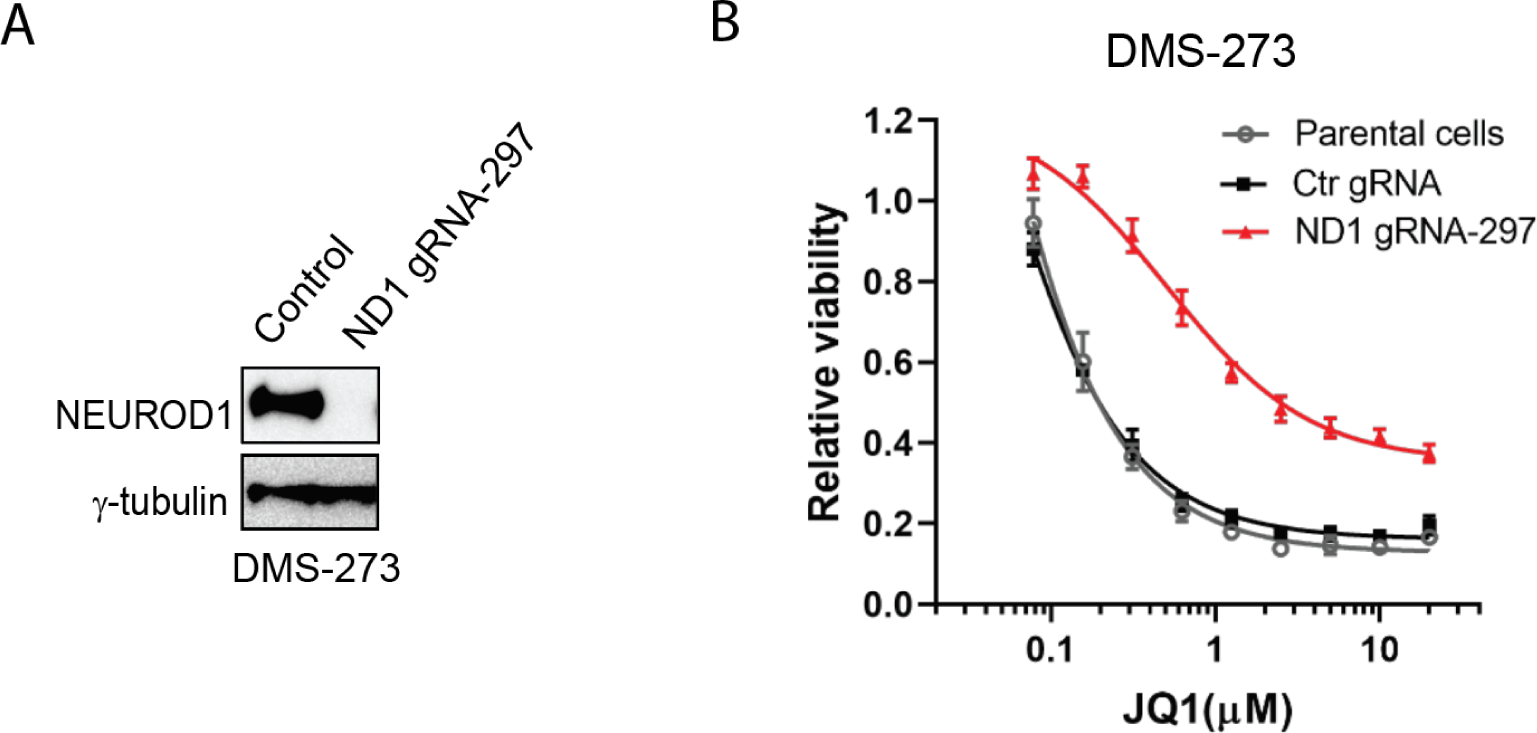
Knockout of NEUROD1 renders DMS-273 cells resistant to BETi. **A)** Immunoblots showing loss of NEUROD1 in DMS-273 cells after CRISPR gene editing with a NEUROD1- targeting gRNA (#297) versus the cells receiving a control gRNA. **B)** Viability of the DMS-273 NEUROD1-KO cells versus the control and parental cells after JQ1 treatment (72 hrs). Error bars represent the SD of four replicates.

**Figure S10.**
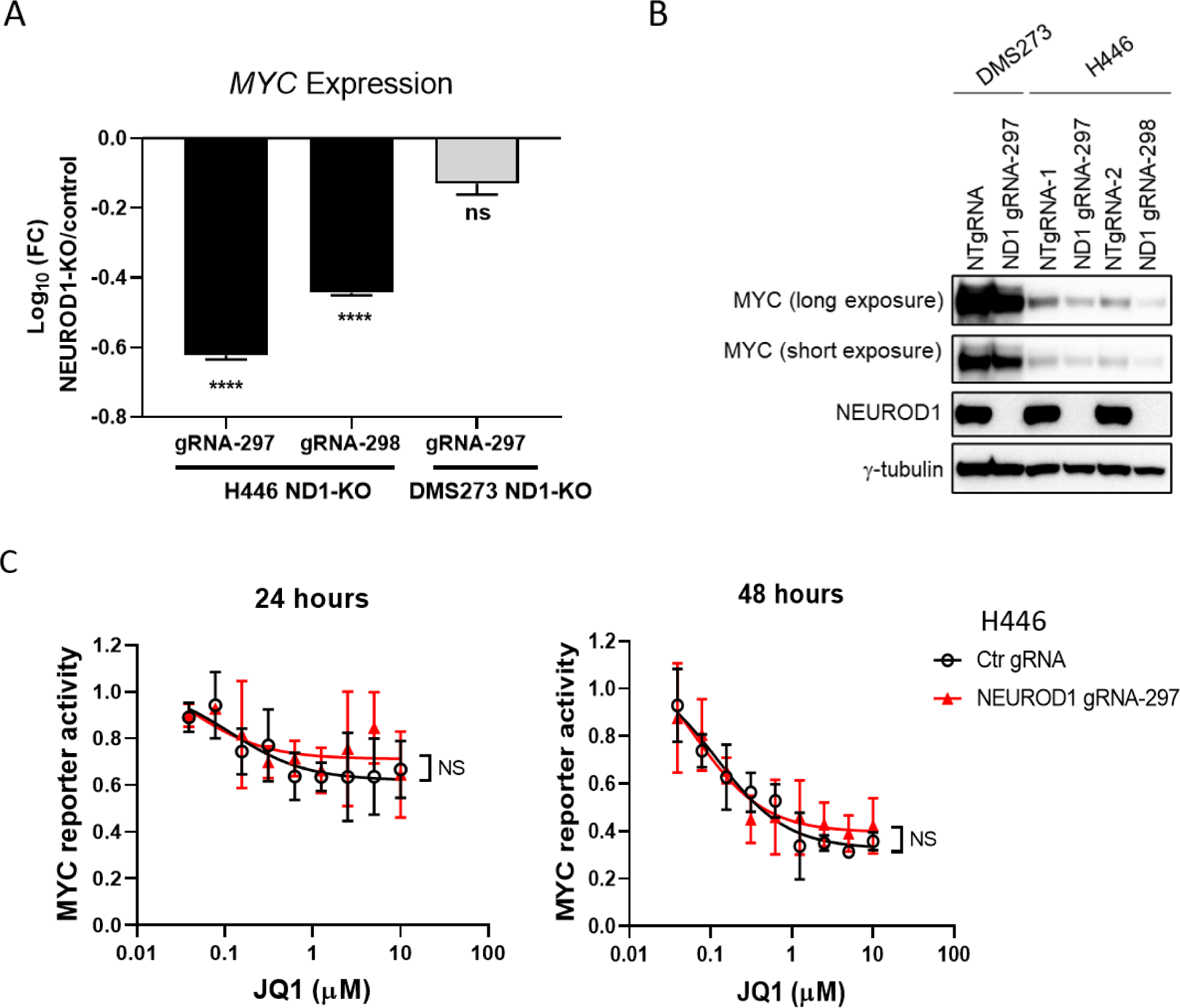
Effects of NEUROD1 KO on MYC expression and transcriptional activity. A-B) NEUROD1 KO modestly decrease of *MYC* gene (A) and protein expression (B) in the H446 cells but not in DMS273 cells. C) JQ1 equally suppressed MYC transcriptional activity in H446 NEUROD1-KO and control cells (left, 24-hr time interval; right, 48-hr time interval). The two- group comparison was performed using the Student’s t-test. NS, no significance.

**Figure S11.**
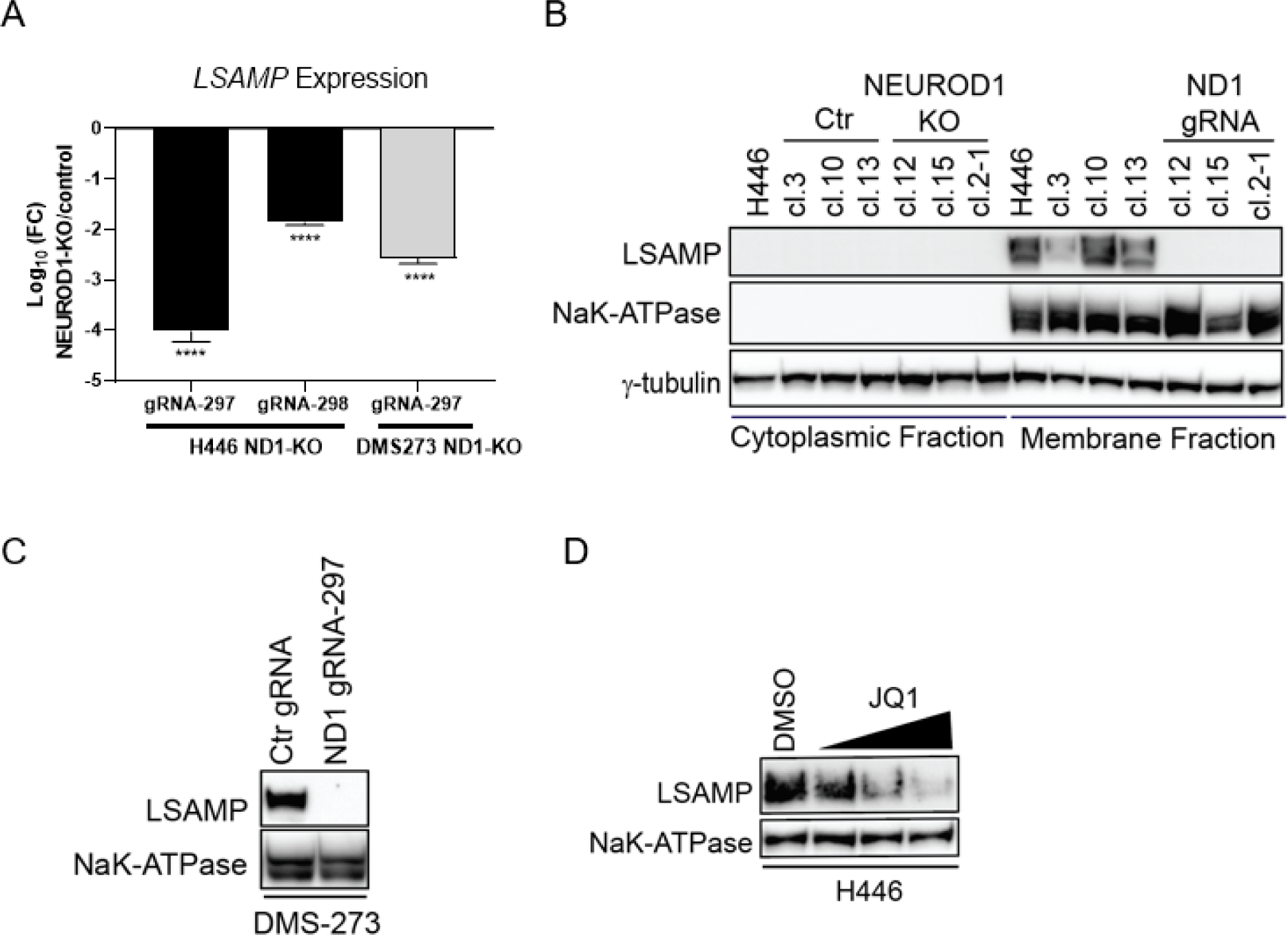
LSAMP expression is regulated by both NEUROD1 and BET bromodomain proteins. **A)** A drastic decrease of *LSAMP* gene expression in the H446 and DMS-273 cells upon KO of NEUROD1. The gene expression was measured by qRT-PCR and plotted relative to control cells. **B)** LSAMP protein is present only in the membrane-bound protein fractions in H446 control cells but not in the NEUROD1-KO cells. **C)** Loss of LSAMP protein expression in DMS273 cells upon KO of NEUROD1. **D)** A dose-dependent decrease of LSAMP protein expression in H446 cells 24 hours after JQ1 treatment (0.25, 0.5, and 1 µM).

**Figure S12.**
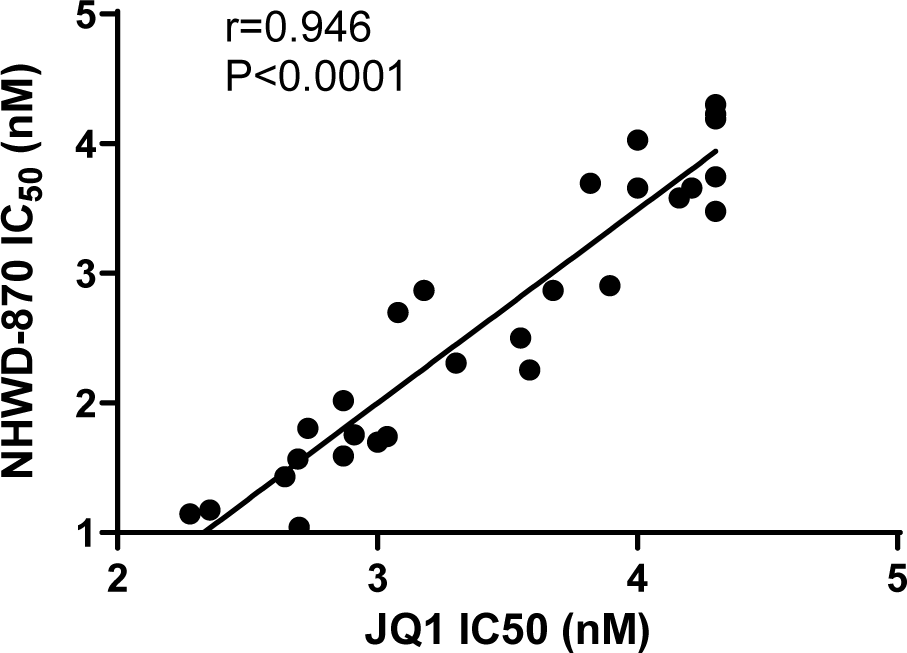
Correlation between the IC50 values of NHWD-870 and JQ1 in SCLC cell lines (N=28). Pearson correlation coefficient and P value were shown.

**Table S1.** The list of the SCLC cell lines (N=52) used in this study.

| Cell Line | Subtype | Source* | Catalog # | Growth media |
| --- | --- | --- | --- | --- |
| COLO-668 | ASCL1 | ATCC | 87061209 | RPMI + 10% FBS |
| COR-L47 | ASCL1 | Sigma-Aldrich | 92031915 | RPMI + 10% FBS |
| COR-L88 | ASCL1 | Sigma-Aldrich | 92031917 | RPMI + 10% FBS |
| DMS454 | ASCL1 | ATCC | 95062832 | Waymouth + 1x glutamax + 10% FBS |
| DMS53 | ASCL1 | ATCC | CRL-2062 | RPMI + 10% FBS |
| DMS79 | ASCL1 | Sigma-Aldrich | 95062824 | RPMI + 10% FBS |
| H1092 | ASCL1 | ATCC | CRL-5855 | HITES<sup>†</sup> + 5% FBS |
| H1105 | ASCL1 | ATCC | CRL-5856 | HITES<sup>†</sup> + 5% FBS |
| H1417 | ASCL1 | ATCC | CRL-5872 | RPMI + 10% FBS |
| H1436 | ASCL1 | ATCC | CRL-5871 | HITES<sup>†</sup> + 5% FBS |
| H146 | ASCL1 | ATCC | HTB-173 | RPMI + 10% FBS |
| H1836 | ASCL1 | ATCC | CRL-5898 | HITES<sup>†</sup> + 5% FBS |
| H187 | ASCL1 | ATCC | CRL-5804 | RPMI + 10% FBS |
| H1876 | ASCL1 | ATCC | CRL-5902 | HITES<sup>†</sup> + 5% FBS |
| H1882 | ASCL1 | ATCC | CRL-5903 | HITES<sup>†</sup> + 5% FBS |
| H1930 | ASCL1 | ATCC | CRL-5906 | RPMI + 10% FBS |
| H1963 | ASCL1 | ATCC | CRL-5982 | RPMI + 10% FBS |
| H2029 | ASCL1 | ATCC | CRL-5913 | HITES<sup>†</sup> + 5% FBS |
| H2081 | ASCL1 | ATCC | CRL-5920 | HITES<sup>†</sup> + 5% FBS |
| H209 | ASCL1 | ATCC | HTB-172 | RPMI + 10% FBS |
| H2107 | ASCL1 | ATCC | CRL-5983 FL | HITES<sup>†</sup> + 5% FBS |
| H2141 | ASCL1 | ATCC | CRL-5927 | HITES<sup>†</sup> + 5% FBS |
| H2196 | ASCL1 | ATCC | CRL-5932 | HITES<sup>†</sup> + 5% FBS |
| H2227 | ASCL1 | ATCC | CRL-5934 | HITES<sup>†</sup> + 5% FBS |
| H345 | ASCL1 | ATCC | HTB-180 | HITES<sup>†</sup> + 5% FBS |
| H378 | ASCL1 | ATCC | CRL-5808 | RPMI + 10% FBS |
| H510A | ASCL1 | ATCC | HTB-184 | F12K + 10% FBS |
| H64 | ASCL1 | ATCC | CRL-5976 | HITES<sup>†</sup> + 5% FBS |
| H69 | ASCL1 | ATCC | HTB-119 | RPMI + 10% FBS |

| Cell Line | Subtype | Source* | Catalog # | Growth media<sup>†</sup> |
| --- | --- | --- | --- | --- |
| H889 | ASCL1 | ATCC | CRL-5817 | RPMI + 10% FBS |
| HCC-33 | ASCL1 | ATCC | ACC 487 | RPMI + 20% FBS |
| COR-L279 | NEUROD1 | Sigma-Aldrich | 96020724 | RPMI + 10% FBS |
| CPC-N | NEUROD1 | DSMZ | ACC 306 | McCoy's 5A + 10% FBS |
| DMS 273 | NEUROD1 | Sigma-Aldrich | 95062830 | Waymouth + 1x glutamax + 10% FBS |
| H1694 | NEUROD1 | ATCC | CRL-5888 | HITES<sup>†</sup> + 5% FBS |
| H2066 | NEUROD1 | ATCC | CRL-5917 | HITES<sup>†</sup> + 5% FBS |
| H2171 | NEUROD1 | ATCC | CRL-5929 | HITES<sup>†</sup> + 5% FBS |
| H446 | NEUROD1 | ATCC | HTB-171 | RPMI + 10% FBS |
| H524 | NEUROD1 | ATCC | CRL-5831 | RPMI + 10% FBS |
| H82 | NEUROD1 | ATCC | HTB-175 | RPMI + 10% FBS |
| SCLC-21H | NEUROD1 | DSMZ | ACC 372 | DMEM + 10% FBS |
| SCLC-22H | NEUROD1 | DSMZ | ACC 373 | DMEM + 10% FBS |
| COR-L311 | POU2F3 | Sigma-Aldrich | 96020721 | RPMI + 10% FBS |
| H1048 | POU2F3 | ATCC | CRL-5853 | HITES<sup>†</sup> + 5% FBS |
| H211 | POU2F3 | ATCC | CRL-5824 | RPMI + 10% FBS |
| H526 | POU2F3 | ATCC | CRL-5811 | RPMI + 10% FBS |
| DMS114 | YAP1 | ATCC | CRL-2066 | Waymouth + 1x glutamax + 10% FBS |
| H1341 | YAP1 | ATCC | CRL-5864 | ACL-4<sup>‡</sup> + 5% FBS |
| H196 | YAP1 | ATCC | CRL-5823 | RPMI + 10% FBS |
| H841 | YAP1 | ATCC | CRL-5845 | HITES<sup>†</sup> + 5% FBS |
| SBC3 | YAP1 | JCRB | JCRB0818 | EMEM + 10% FBS |
| SBC5 | YAP1 | JCRB | JCRB0819 | EMEM + 10% FBS |

\*ATCC, American Type Culture Collection; DSMZ, German Collection of Microorganisms and Cell Cultures; JCRB, Japanese Collection of Research Bioresources Cell Bank.
<sup>†</sup> HITES contains DMEM:F12 medium (ATCC, #30-2006; 500 ml), Insulin (Sigma, #I1882; 0.005mg/ml), apo-transferrin (Sigma, #T5391 or T1428; 0.01 mg/ml), sodium selenite (Sigma, #S9133; 30 nM), hydrocortisone (Sigma, #H0135; 10nM), $\beta$ -estradiol (Sigma, #E2257; 10nM), glutamax (Gibco, #35050-061; 1x).
<sup>‡</sup> ACL4 contains DMEM:F12 medium (ATCC, #30-2006; 500 ml), insulin (Sigma, #I1882; 0.02mg/ml), apo-transferrin (Sigma, #T5391 or T1428; 0.01 mg/ml), sodium selenite (Sigma, #S9133; 25 nM), hydrocortisone (Sigma, #H0135; 50nM), EGF (Sigma, #SRP3027; 1ng/ml), glutamax (Gibco, #35050-061; 1x), ethanolamine (Sigma, #E0135; 0.01mM), phosphorylethanolamine (Sigma, #P0503; 0.01mM), triiodothyronine (Sigma, #T6397; 100pM), HEPES (Sigma, #H0887; 10mM), sodium pyruvate (Gibco, #11360-070; 0.5mM).

**Table S2.**
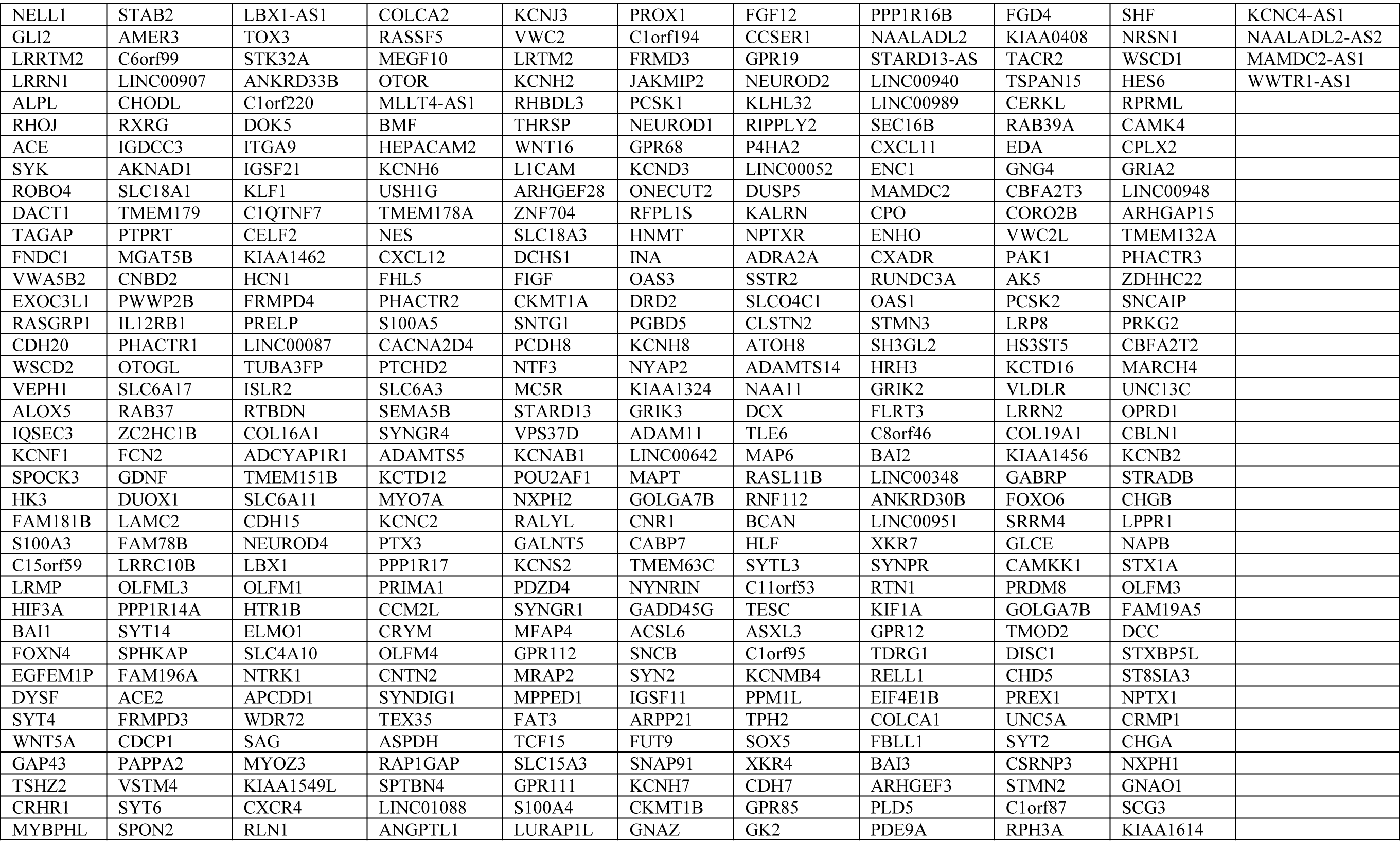
The list of genes included in the NEUROD1 gene signature.

**Table S3.**
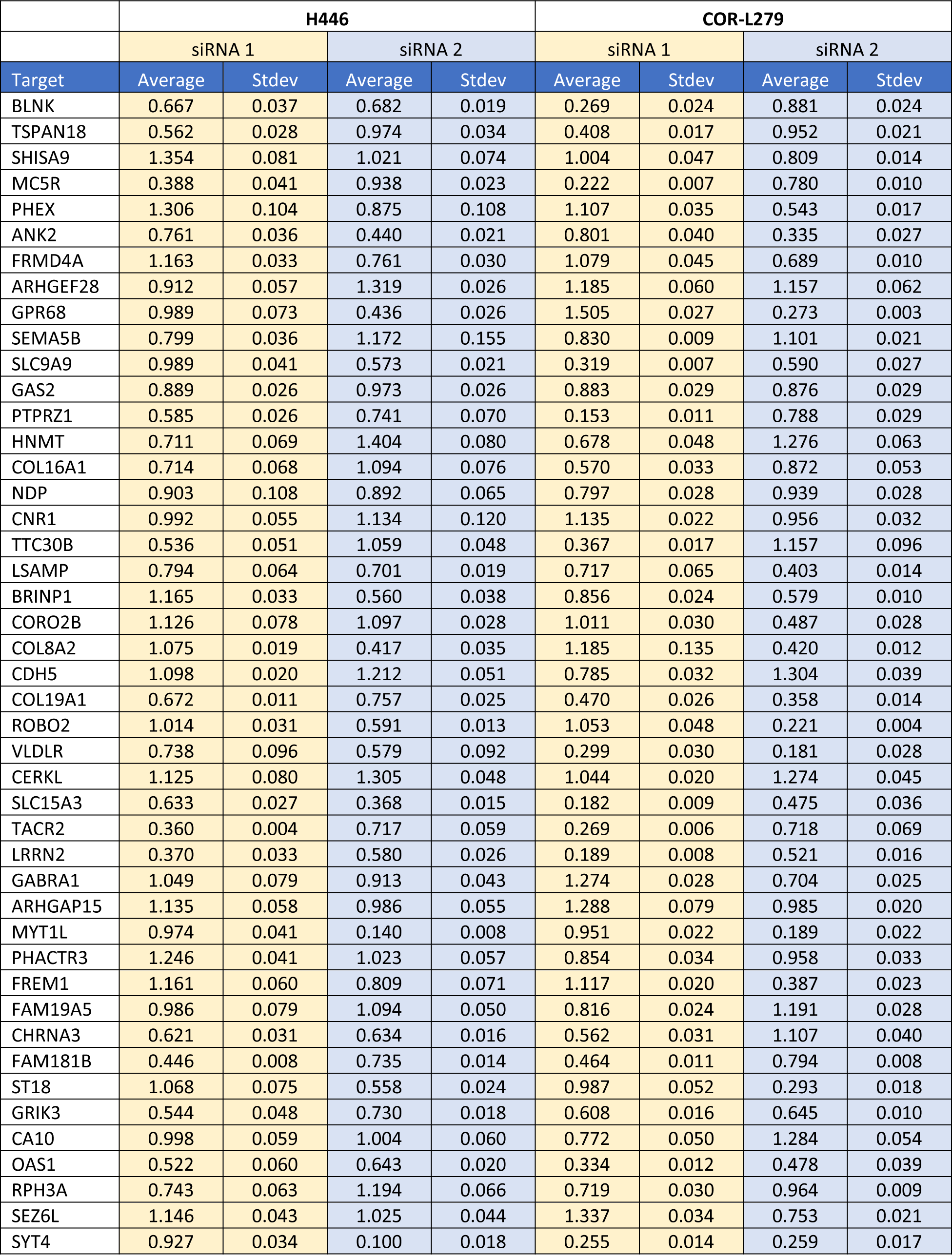
The siRNA screen results in H446 and COR-L279 cells.

